# Mechanism of beta-arrestin 1 mediated Src activation via Src SH3 domain revealed by cryo-electron microscopy

**DOI:** 10.1101/2024.07.31.605623

**Authors:** Natalia Pakharukova, Brittany N. Thomas, Harsh Bansia, Linus Li, Dana K. Bassford, Rinat R. Abzalimov, Jihee Kim, Alem W. Kahsai, Biswaranjan Pani, Kunhong Xiao, Roni Ochakovski, Shibo Liu, Xingdong Zhang, Seungkirl Ahn, Amedee des Georges, Robert J. Lefkowitz

**Affiliations:** Department of Medicine, Duke University Medical Center; Durham, NC 27710, USA; Howard Hughes Medical Institute, Duke University Medical Center; Durham, NC 27710, USA; Department of Molecular Pathobiology, College of Dentistry, New York University, New York, 10010, USA; Pain Research Center, New York University, New York, 10010, USA; Structural Biology Initiative, CUNY Advanced Science Research Center; New York, NY 10031, USA; Center for Proteomics & Artificial Intelligence, Allegheny Health Network Cancer Institute, Pittsburgh, PA 15202, USA; Department of Biomedical Engineering, College of Engineering, Carnegie Mellon University, Pittsburgh, PA 15213, USA; Department of Biochemistry, Duke University Medical Center; Durham, NC 27710, USA

## Abstract

Beta-arrestins (βarrs) are key regulators and transducers of G-protein coupled receptor signaling; however, little is known of how βarrs communicate with their downstream effectors. Here, we report the first structural insights into the fundamental mechanisms driving βarr-mediated signal transduction. Using cryo-electron microscopy, we elucidate how βarr1 recruits and activates the non-receptor tyrosine kinase Src, the first identified signaling partner of βarrs. βarr1 engages Src SH3 through two distinct sites, each employing a different recognition mechanism: a polyproline motif in the N-domain and a non-proline-based interaction in the central crest region. At both sites βarr1 interacts with the aromatic surface of SH3, disrupting the autoinhibited conformation of Src and directly triggering its allosteric activation. This structural evidence establishes βarr1 as an active regulatory protein rather than a passive scaffold and suggests a potentially general mechanism for βarr-mediated signaling across diverse effectors.

## Main text

Beta-arrestins 1 and 2 (βarr1 and βarr2) are ubiquitously expressed scaffold proteins that interact with most, if not all, G-protein coupled receptors (GPCRs)^1^. βarrs mediate receptor desensitization and intracellular trafficking, as well as initiate downstream signaling cascades, independently or in concert with G-proteins^2^. In their role as signal transducers, βarrs link GPCRs with numerous downstream effectors, including components of several mitogen-activated protein kinase cascades and Src family kinases, amongst many others^2^. Whereas early studies suggested that βarrs function as signaling scaffolds, recent findings revealed that βarrs also directly allosterically activate some of their effectors such as non-receptor tyrosine kinase Src^3, 4^, mitogen-activated protein kinases c-Raf^5^, and extracellular signal-regulated kinase 2^6^. However, the molecular mechanisms of βarr-mediated signaling remain elusive, largely due to the absence of any structural data on complexes of βarrs with any of their diverse effector enzymes. The transient nature of βarr–effector complexes and the mid-micromolar affinity of their interactions markedly complicates their structural studies, necessitating the use of stabilization techniques.

Src homology 3 (SH3) domains are one of the most ubiquitous binding modules with nearly 300 members found in the human genome^7^. SH3-containing proteins are involved in various signaling pathways, such as cell growth and proliferation, endocytosis, and cytoskeleton remodeling^8^. Notably, βarrs have been shown to interact with the SH3 domains of several proteins^3^, including the proto-oncogene kinase Src, the first discovered effector protein of βarrs^9^. Intriguingly, βarrs lack the canonical left-handed type II polyproline motifs required for SH3 binding; therefore, how βarrs recruit SH3 remains unknown. Here, we report the first structural insights into the fundamental mechanisms of signal transduction by βarrs. Using Src as a prototypical effector of βarr, we elucidate the molecular basis for Src SH3 recruitment and Src allosteric activation by cryo-electron microscopy (cryo-EM).

## Results

### βarr1 uses two distinct sites to bind SH3

To map the binding interface between the SH3 domain of Src and βarr1, we utilized a disulfide trapping strategy^10^. The βarr1 and SH3 Src sequences were derived from rat (*Rattus norvegicus*) and chicken (*Gallus gallus*), respectively. Both proteins exhibit high inter-species sequence similarity—93.54% for βarr1 and 93.66% for Src (100% for Src SH3 domain), with the most notable sequence variations present in the unique domain of Src and at the C-terminus of βarr1 (Supplementary Fig. 1). We introduced cysteine substitutions at 18 different positions in βarr1 in proximity to P88-P91 and P120-P124, previously reported to be critical for βarr1–Src interaction (Fig. 1a)^3, 9^. Guided by the available structures of SH3 domains in complex with polyproline-rich peptides, we designed 13 cysteine mutants of SH3 (Fig. 1b). We comprehensively tested 234 combinations of purified βarr1 and SH3 cysteine mutant pairs by inducing disulfide bond formation with hydrogen peroxide *in vitro*. Active βarr1 was shown to bind more strongly to SH3^4^; therefore, prior to disulfide trapping reactions βarr1 was activated using a synthetic phosphopeptide mimicking the C-tail of vasopressin 2 receptor (V2Rpp) and the stabilizing antibody fragment Fab30. Intriguingly, we observed the formation of disulfide cross-linked SH3– βarr1–V2Rpp–Fab30 complexes by two βarr1 mutants, E92C in the distal part of the N-domain (hereafter referred to as **βarr1-N** site) and P120C in the central crest region (hereafter referred to as **βarr1-CC** site) (Fig. 1c, d). In contrast, little to no SH3–βarr1 cross-linked complexes were formed by other βarr1 mutants, such as T116C (Fig. 1d). The SH3 mutant R95C in the RT-loop gave the strongest disulfide cross-linked band in both sites. As βarr1-N and βarr1-CC sites are more than 20 Å apart, we hypothesized that the SH3 domain might independently bind at two sites of βarr1. Complex formation at the βarr1-CC site was more efficient in the presence of V2Rpp and Fab30, whereas the βarr1-N site forms the complex equally well regardless of activation (Supplementary Fig. 2a).

**Fig. 1.**
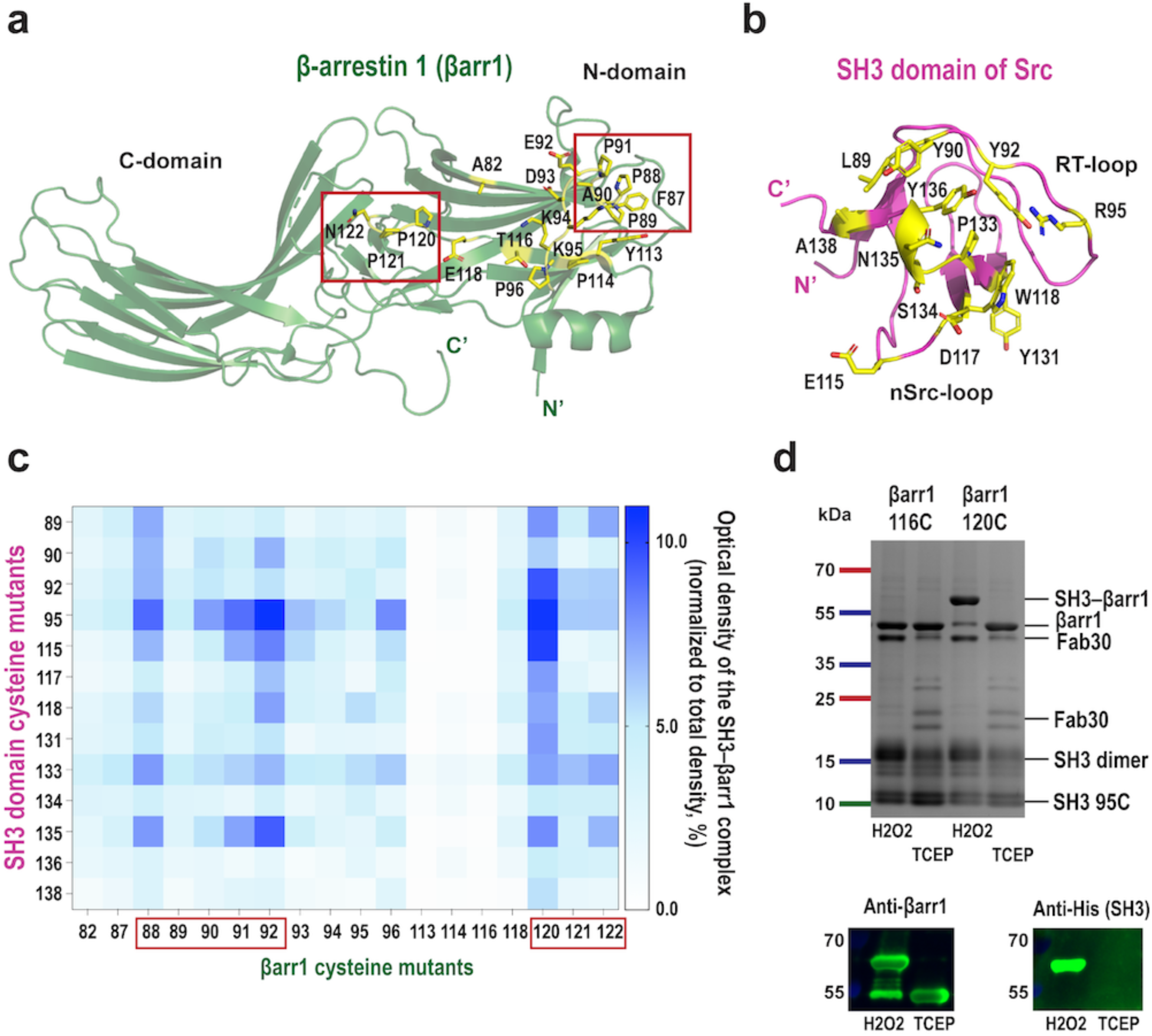
βarr1 uses two distinct sites to bind SH3. **a, b** Structures of βarr1 (green, PDB: 4JQI) and SH3 (magenta, PDB: 2PTK), cartoon representation. Residues used for disulfide trapping are shown as yellow sticks and labeled. Proline regions of βarr1 are framed in rectangles. **c,** Heatmap of different SH3–βarr1 cysteine-pair complexes; densitometry analysis of Coomassie blue gels. The SH3–βarr1 complex band was normalized to the total density (mean; n=3-4). **d,** Top: representative results of disulfide trapping with βarr1_120C (binding site for SH3) and βarr1_116C (non-binding site for SH3) (Coomassie blue gel). Bottom: Western blot of disulfide trapping reaction between βarr1_120C and SH3_R95C confirms that the 60-kDa band is the SH3–βarr1 complex.

SH3 E115C also formed a significant amount of cross-linked complex with βarr1-CC (Fig. 1c). Analytical size-exclusion chromatography revealed that the SH3_R95C–βarr1_P120C complex displays a more prominent and symmetric peak compared to SH3_E115C–βarr1_P120C, suggesting greater homogeneity and stability (Supplementary Fig. 2b). βarr1 mutants P88C and P91C mutants also formed the cross-linked complex with SH3_R95C, but to a lesser degree than E92C suggesting the dynamic nature of the SH3–βarr1 interfaces (Fig. 1c).

To validate whether V2Rpp-activated βarr1 can bind SH3 at two distinct sites, we used isothermal titration calorimetry (Supplementary Fig. 2c). The integrated heat curve revealed two binding events: one with a Kd of ∼6 μM and the other with a significantly lower affinity of ∼50 μM. The favorable entropy (-TΔS) suggests that in both sites, SH3–hydrophobic interactions drive βarr1 binding. Furthermore, the favorable enthalpy (ΔH) at the higher affinity site indicates some contribution from hydrogen bonding and van der Waals forces.

### βarr1 binds to the aromatic surface of SH3

To elucidate the mechanism of SH3 recruitment by βarr1, we formed cross-linked complexes with the best mutant pairs SH3_95C–βarr1_120C (**SH3–βarr1-CC**) and SH3_95C–βarr1_92C (**SH3– βarr1-N**) and determined the structures by cryo-EM (Fig. 2a-c, Supplementary Fig. 3, Supplementary Fig. 4, Supplementary Table 1). The map obtained from the SH3–βarr1-CC complex has a global resolution of 3.47 Å and showed a rigid interaction with SH3 in the central crest area of βarr1 (Fig. 2a, Supplementary Fig. 4e). The complex was activated by V2Rpp and stabilized by Fab30 and nanobody 32 (Nb32)^11^. βarr1 adopts an active conformation^12^, characterized by C-tail displacement and inter-domain rotation, and binds to the aromatic surface of SH3 containing residues from the RT loop, nSrc loop, and the 3_10_ helix (Fig. 2b, Supplementary Video 1). βarr1 interacts with SH3 using β-strand V (residues 75-80) in the N domain and a part of the lariat loop in the C domain. Y90 and Y136 in SH3 interact with F75 in βarr1. The side chains of Y136 and N135 in SH3 are well-placed to make hydrogen bonds with the main chain oxygen of R76 in βarr1. Additional hydrogen bonds are possible between the main chain oxygen of E93 in SH3 and the side chain of N122 in βarr1, as well as between W118 in SH3 and the main chain oxygen of D78 in βarr1.

**Fig. 2.**
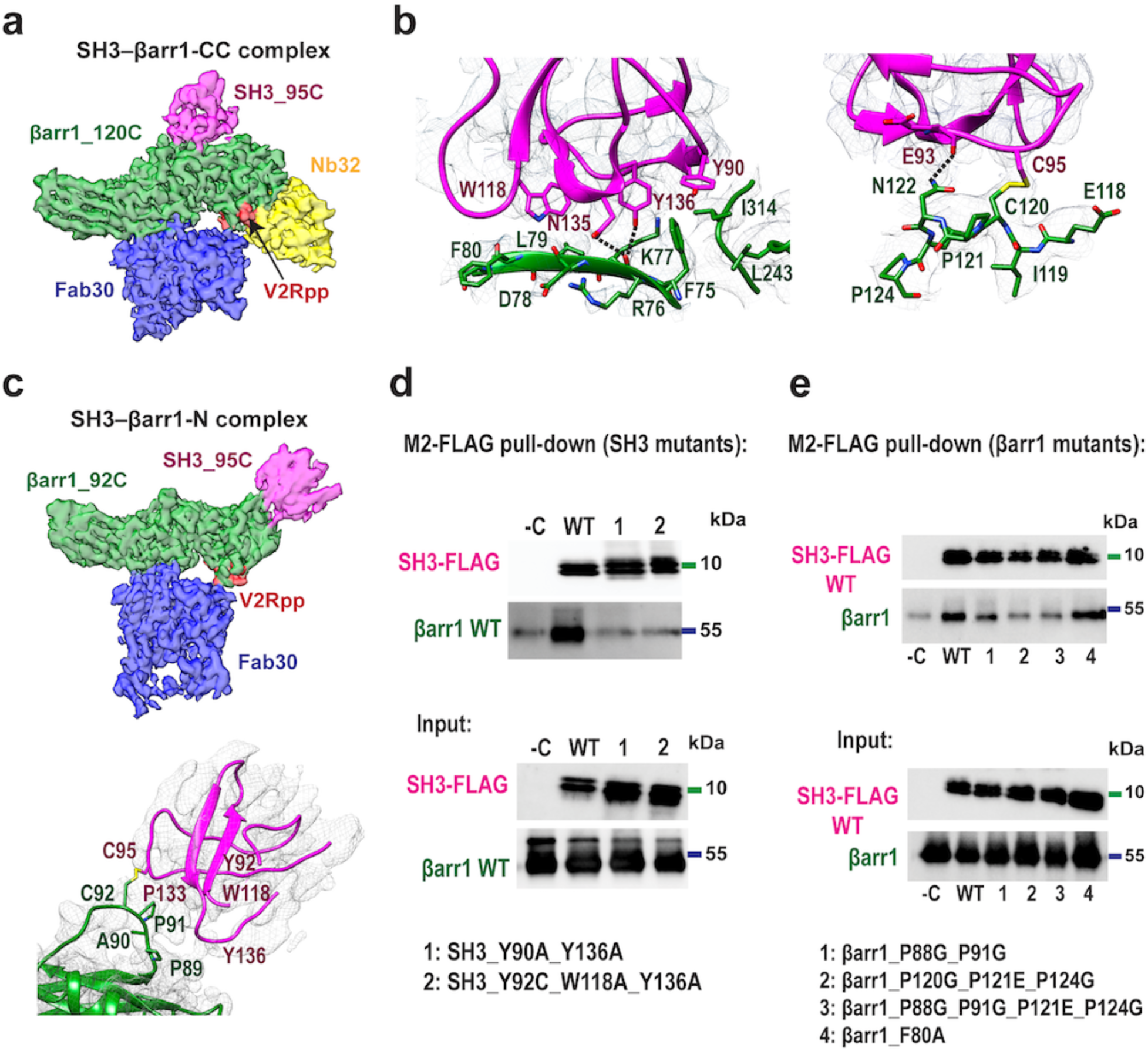
βarr1 binds the aromatic surface of SH3. **a,** Cryo-EM density map of the SH3–βarr1-CC complex (green, βarr1; magenta, SH3; red, V2Rpp; blue, Fab30; yellow, Nb32). Map contour level is 0.75. **b,** Interaction interface of SH3–βarr1-CC complex (green, βarr1; magenta, SH3) with density mesh (upsample map 0.69 Å/pix, contour level 0.37). Hydrogen bonds are shown as dashed lines. See also Supplementary Video 1. **c,** Top panel: Cryo-EM density map of the SH3–βarr1-N complex (green, βarr1; magenta, SH3; red, V2Rpp; blue, Fab30). Map contour level is 0.50. Bottom panel: The interaction interface of SH3–βarr1-N complex (green, βarr1; magenta, SH3) with density mesh (upsample map 0.69 Å/pix, contour level 0.2). Only the backbone of SH3 was modeled; βarr1 side chains are shown for illustrative purposes. See also Supplementary Video 2. **d,** Representative Western blots of M2-FLAG pull-down assay of SH3-FLAG mutants and βarr1 WT (n=3). **e,** Representative Western blots of M2-FLAG pull-down assay of SH3-FLAG WT and βarr1 mutants (n=3).

The SH3–βarr1-N complex was formed with V2Rpp and Fab30 but without Nb32, as Nb32 hinders complex formation due to the steric clashes with SH3. The cryo-EM map showed SH3 bound to the distal part of the N-terminal domain of βarr1 (Fig. 2c). Whereas SH3–βarr1-CC has a clearly resolved density for SH3, in SH3–βarr1-N, the density for SH3 is weaker (local resolution is 4.2-4.6 Å) since SH3 binds to a flexible loop of βarr1, the ^87^F<u>PPAP</u>EDK^94^ motif (Fig. 2c, Supplementary Fig. 4f). We used molecular dynamics flexible fitting (MDFF) constrained by the position of the disulfide bond between βarr1 and SH3 to place the main chain of the SH3 domain. The proline region of the βarr1-N site (P88, P89, P91) appears to interact with the aromatic surface of SH3, similarly to the βarr1-CC site (Fig. 2c, Supplementary Video 2). However, the weak density for SH3 in the SH3–βarr1-N complex prevents confident identification of the specific SH3 residues involved in βarr1 binding. To confirm the interaction interfaces, we introduced mutations in SH3 and βarr1 and tested the binding by pull-down assay. Mutations in the RT loop (Y90A, Y92C) and the 3₁₀ helix (Y136A) of SH3 drastically reduced its ability to bind βarr1 (Fig. 2d). Likewise, the substitutions of prolines significantly impair SH3 binding (Fig. 2e). These findings, together with the disulfide trapping data indicating that the RT loop faces the SH3–βarr1 interaction interface, confirm that residues within both the RT loop and 3₁₀ helix are critical for the SH3–βarr1 interaction at both sites of βarr1.

The structural and disulfide trapping data indicate a highly dynamic mode of SH3 binding to the βarr1-N site (Fig. 1c), with disulfide trapping capturing only a single state of this inherently dynamic interaction. Cross-linked complex formation by several SH3–βarr1 cysteine pairs (Fig. 1c) suggests that SH3 might adopt alternative orientations relative to the βarr1-N site, highlighting the complexity of low-affinity interactions in signaling cascades.

Notably, βarr1 uses a distinct recognition mechanism for SH3 binding at each site. The βarr1-N site binds SH3 exclusively via proline residues. In contrast, the recruitment of SH3 via the βarr1-CC site is driven by the non-proline residues in the β-strand V and the lariat loop and two proline residues after the β-strand VI.

Since the SH3–βarr1 complexes were stabilized using cysteine substitutions and disulfide trapping, we sought to determine whether these interactions also occur under physiological conditions, in the absence of engineered mutations. To address this, we assessed the solvent accessibility of free βarr1, V2Rpp-activated βarr1, and SH3 using hydrogen–deuterium exchange mass spectrometry (HDX-MS) (Fig. 3). HDX-MS provides insights into protein interaction interfaces by measuring deuterium uptake in peptides typically 5–15 residues in length. Thus, this approach enables us to verify whether SH3 binds to two distinct regions of βarr1 *in vitro* without cross-linking. Consistent with the structures and the pull-down data, βarr1 showed a significant decrease in HDX rate in both the βarr1-CC and βarr1-N sites in the presence of SH3, and this effect was more prominent for V2Rpp-activated βarr1 (Fig. 3). The nSrc loop and the 3_10_ helix of SH3 also demonstrated a decrease in the HDX uptake (Supplementary Fig. 5). Taken together with the structural data and disulfide trapping results, these findings support that the SH3–βarr1-CC and SH3–βarr1-N structures represent *bona fide* interaction complexes.

**Fig. 3.**
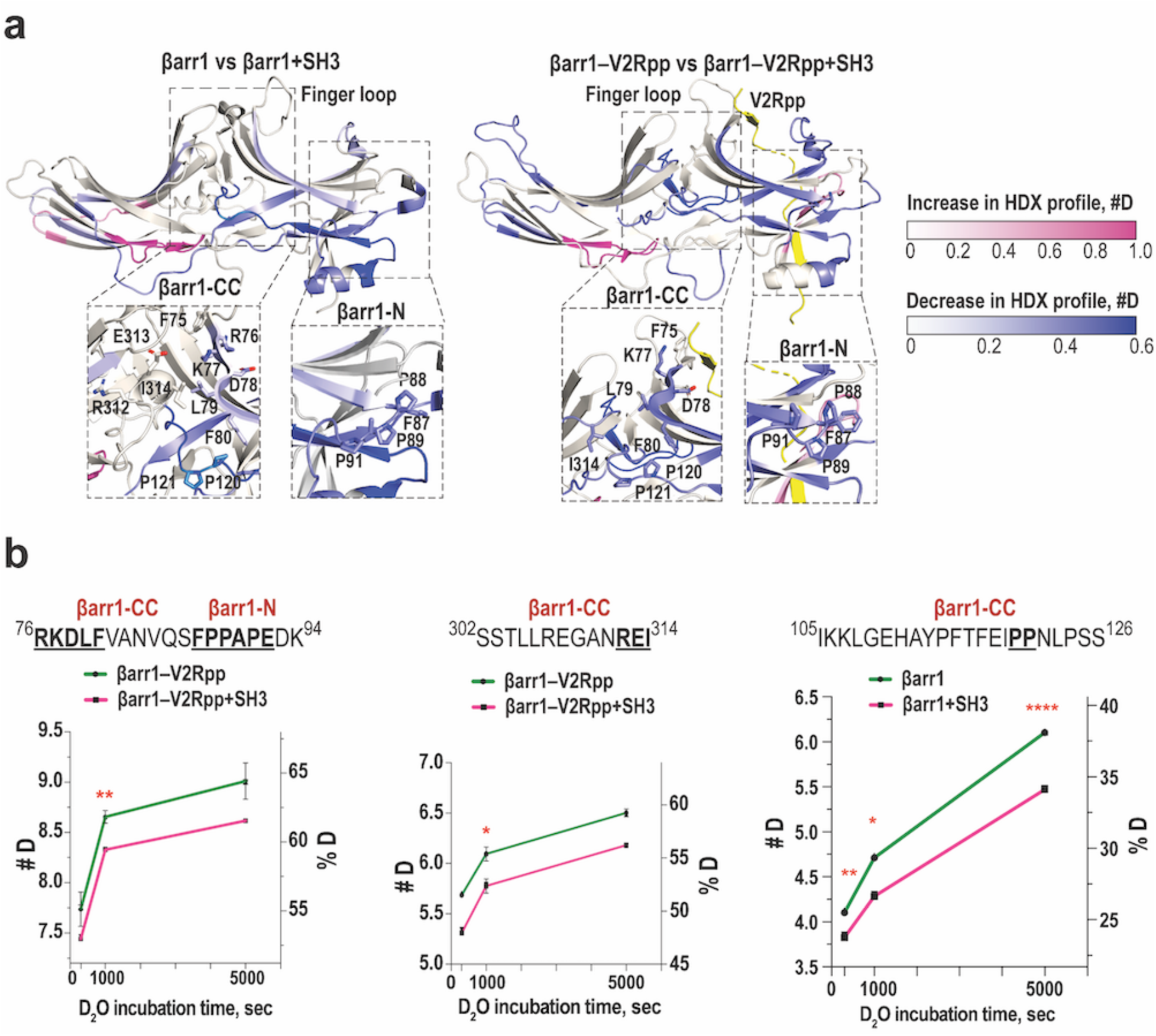
HDX profile changes of free and V2Rpp-activated βarr1 in the presence of SH3. **a,** Structures of free βarr1 (PDB: 1G4M) and V2Rpp-activated βarr1 (PDB: 4JQI, βarr1 - grey; V2Rpp - yellow); regions with decreased and increased deuterium uptake are shaded in purple and pink, respectively. Insets: focus on the βarr1-N and βarr1-CC sites. **b,** The HDX changes of βarr1 peptides corresponding to βarr1-CC and βarr1-N sites upon co-incubation with SH3 (mean ± SEM, n=3). The residues interacting with SH3 in the structures are shown in bold and underlined. Statistical analysis was performed using Student’s t-test (*p < 0.05, **p < 0.01, ****p < 0.0001).

We then compared the mechanisms of SH3 recruitment by various SH3-binding proteins and βarr1-N (Supplementary Fig. 6a-c) and βarr1-CC sites (Supplementary Fig. 6d, e)^13–17^. In all structures, SH3 utilizes its aromatic surface to engage with its binding partners (Supplementary Fig. 6). The βarr1-N site resembles the canonical polyproline-rich motif observed in Nef and ELMO1, although the polyproline sequence in βarr1 is considerably shorter (Supplementary Fig. 6a, c). In contrast, the βarr1-CC site employs non-proline residues to recruit SH3 (Fig. 2b, Supplementary Fig. 6d, e). Non-canonical binding was previously reported for several SH3-binding proteins^18, 19^. For example, endophilin A1 SH3 interacts with histidine, proline and two arginine residues in parkin Ubl (Supplementary Fig. 6b); SH3 of Sla1 endocytic protein engages with histidine and hydrophobic residues of ubiquitin (Supplementary Fig. 6e)^15, 16^. However, βarr1 uniquely employs two distinct sites and two different recognition mechanisms to bind SH3.

### βarr1 drives autoinhibition relief of Src

To understand how βarr1 activates Src, we aimed to determine the structure of βarr1 in complex with three-domain Src (SH3-SH2-SH1, residues 83-533). We performed disulfide trapping experiments using the most promising βarr1 and Src mutants, based on the results of the βarr1– SH3 screening (Fig. 1c, d). Consistent with the findings for the isolated SH3 domain, we observed the formation of Src–βarr1 complexes at both βarr1-N and βarr1-CC sites, with βarr1 P120C (βarr1-CC) forming the highest proportion of the cross-linked complex with Src R95C (Supplementary Fig. 7a, b). Therefore, we formed the Src_R95C–βarr1_P120C complex (hereafter referred to as **Src–βarr1-CC**) and determined its structure by cryo-EM (Fig. 4, Supplementary Fig. 7c, Supplementary Table 1). The final cryo-EM map has a resolution of 3.34 Å with a clear density for the SH3 domain (Fig. 4a, Supplementary Fig. 8a-c). The superposition of SH3–βarr1-CC and the Src–βarr1-CC revealed a translational shift of 2.0-2.2 Å of the SH3 domain towards the central crest of βarr1 in the Src–βarr1-CC complex (Supplementary Fig. 8d). The orientation of SH3 in Src–βarr1-CC and the interaction network with βarr1 are similar to the SH3–βarr1-CC complex (Fig. 4b, Supplementary Fig. 8d). The 2D classes of the complex show fuzzy density around SH3 suggesting the high flexibility of the SH2 and SH1 domains (Fig. 4c). Consistent with this observation, SH2 and SH1 were not resolved in the final map. Filtering the map to a lower resolution (15 Å) revealed the density attributable to SH2 but not to SH1 (Fig. 4d), suggesting that βarr1-CC binding to SH3 induces displacement of the SH1 domain. To determine whether a similar mechanism operates at the βarr1-N site, we formed a complex between Src R95C and βarr1 E92C (**Src–βarr1-N**) and collected a cryo-EM dataset. This dataset had too few usable particles to obtain a reliable 3D reconstruction. Still, using the SH3–βarr1-N map as a reference, we aligned 2D class averages of the Src–βarr1-N complex with projections of the SH3–βarr1-N map (Supplementary Fig. 8e). This comparison revealed that the 2D projections of Src–βarr1-N closely resemble those of the SH3–βarr1-N complex, with the addition of fuzzy densities likely corresponding to the liberated SH2 and SH1 domains, mirroring the SH1 displacement observed in the Src–βarr1-CC complex.

**Fig. 4.**
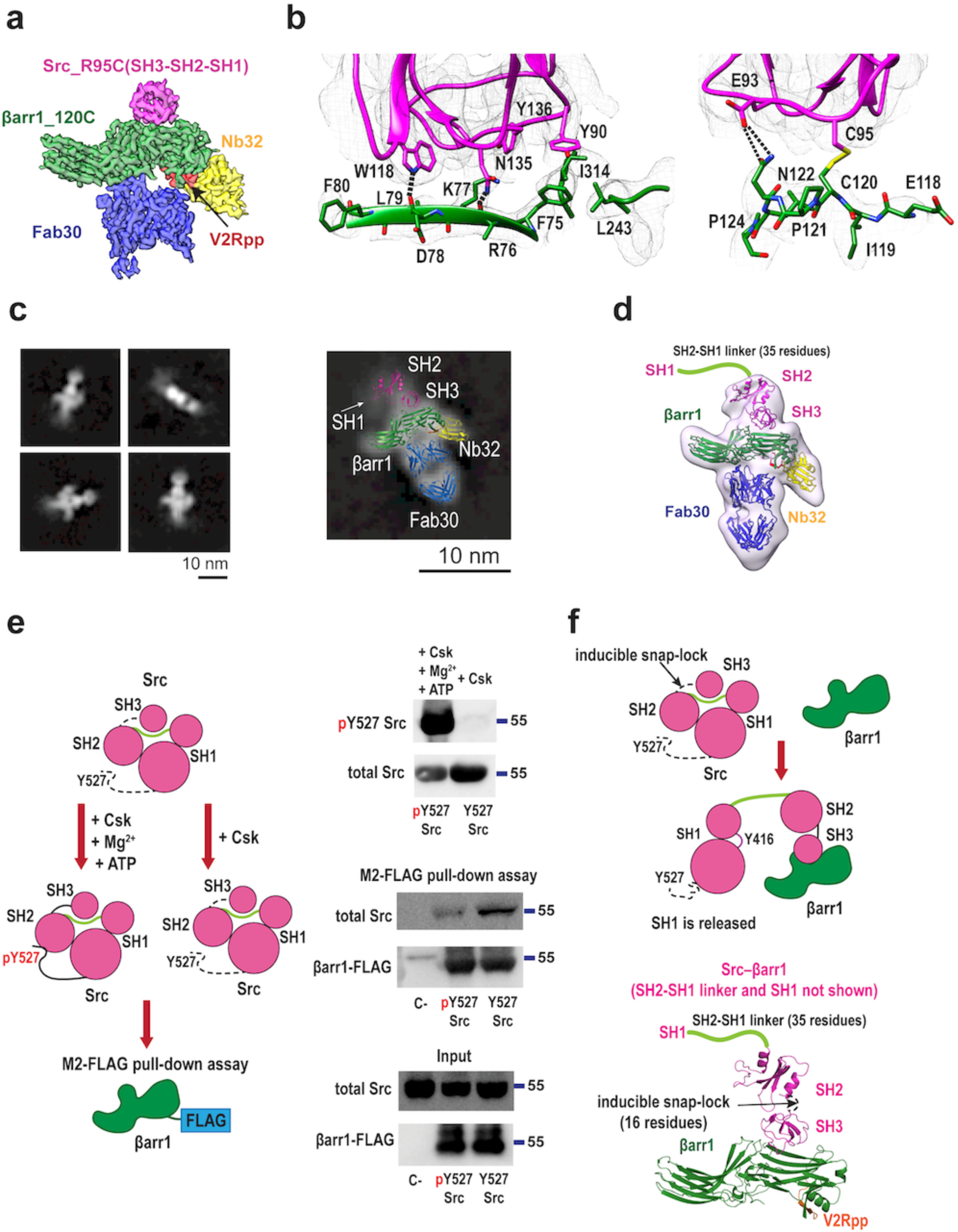
βarr1 drives autoinhibition relief of Src. **a,** Cryo-EM density map of the Src–βarr1-CC complex (green, βarr1; magenta, Src(SH3); red, V2Rpp; blue, Fab30; yellow, Nb32). Map contour level is 0.65. **b**, The interaction interface of Src–βarr1-CC (green, βarr1; magenta, Src (SH3 is shown) with density mesh (upsample map 0.72 Å/pix, contour level 0.15). The hydrogen bonds are shown as dashed lines. See also Supplementary Fig. 8. **c**, Representative 2D classes of Src– βarr1-CC (left) and the 2D class with the fitted model (right). **d,** Cryo-EM density map of the Src– βarr1-CC complex low-pass filtered to 15 Å with the fitted model. Map contour level is 0.06. **e**, βarr1 preferentially binds Src lacking C-tail phosphorylation at Y527. Left panel: schematic of the assay. Right panel, top: representative Western blot of Src phosphorylation by Csk (n=4). Right panel, bottom: representative Western blot of M2-FLAG pull-down assay of βarr1-FLAG and C-tail phosphorylated and C-tail unphosphorylated Src (n=8). **f,** Schematic of βarr1-mediated activation of Src. See also Supplementary Video 3.

Src activity is tightly regulated by three mechanisms of intramolecular interactions named the latch, clamp, and switch that conformationally restrict SH1, the catalytic domain of Src, and maintain its autoinhibited conformation (Supplementary Fig. 9a-c)^20, 21^. Unlatching, unclamping, and switching are required for full activation of Src. The latch involves phosphorylation of C-terminal tail at Y527 and its binding to SH2 (Supplementary Fig. 9b); the clamp mediates binding of SH3 RT loop, the 3_10_ helix and the nSrc loop to the SH2-SH1 linker and SH1 (Supplementary Fig. 9c); the switch involves the autophosphorylation of Src activation loop Y416 by another Src molecule^21^. Interestingly, superimposing the SH3–βarr1 complexes onto the crystal structure of autoinhibited Src phosphorylated at Y527 reveals steric clashes of βarr1 with the SH2 and SH1 domains, suggesting that βarr1 cannot access the SH3 domain in this conformation (Supplementary Fig. 9d). Under physiological conditions, however, the interactions between SH3, SH2, and SH1 domains are thought to be in a dynamic equilibrium between bound and unbound states, allowing the SH3 domain engaged with the SH1-SH2 linker to transiently expose its binding interface to βarr1. The affinities of these interactions are evolutionarily fine-tuned: strong enough to maintain the kinase in an autoinhibited conformation, yet sufficiently weak to respond to diverse cellular stimuli^22^. A conformation of Src lacking C-terminal phosphorylation at Y527 may further favor βarr1 accessibility, as the unphosphorylated tail forms fewer stabilizing interactions with the SH2 domain (Supplementary Fig. 9b). Supporting this, the crystal structure of unphosphorylated Src (PDB: 1Y57) shows the C-tail in a liberated state, with the C-tail-binding pocket on SH2 occupied by a sulfate ion from the crystallization buffer containing 1.3 M ammonium sulfate (Supplementary Fig. 9e). *In vivo*, however, unphosphorylated Src likely exists in a dynamic equilibrium between C-tail-bound and C-tail-liberated states. To determine which Src conformation is favored by V2Rpp-activated βarr1, we compared the binding of βarr1 to Src with and without C-terminal phosphorylation by pull-down assay. Phosphorylation at Y527 was performed *in vitro* using purified Csk, a known negative regulator of Src activity, and confirmed by western blotting (Fig. 4e). Although βarr1 is still capable of binding Y527-phosphorylated Src, it shows stronger binding to the unphosphorylated form, indicating a preference for the more dynamic, flexible conformation of Src (Fig. 4e). Previous studies have shown that the absence of C-tail phosphorylation increases the flexibility of the linker between the SH3 and SH2 domains, known as an “inducible snap lock”, that may further enable the SH3 domain to accommodate not only extended polyproline sequences but also structurally diverse interaction partners^23^. Therefore, in the unphosphorylated state, enhanced flexibility in the SH3–SH2 linker appears to promote βarr1 accessibility by transiently exposing SH3 binding interface to βarr1 (Fig. 4f). As evident from the structures, βarr1 binding to SH3 RT loop and the 3_10_ helix disrupts the clamp mechanism of Src autoinhibition and releases the SH2-SH1 linker and the SH1 domain (Fig. 2a, b, Fig. 4b, Supplementary Fig. 9a). This is further attested by the Src–βarr1-CC structure, in which high flexibility of SH1 is consistent with its unrestricted movement and the disruption of the autoinhibited state. Therefore, βarr1 causes the autoinhibition relief of Src by interacting with SH3 and disrupting the intramolecular clamp (Supplementary Video 3).

In addition to SH3, βarr1 is also known to bind the SH1 domain of Src; however, this interaction does not lead to Src activation^4^. Furthermore, in contrast to SH3, which preferentially interacts with active βarr1, SH1 shows stronger binding to the inactive form^4^. Consistent with this, our structural data show that SH1 is displaced from its position in inactive Src and does not appear to associate with V2Rpp-activated βarr1 in the conditions of our cryo-EM experiments. To gain insight into how inactive βarr1 engages SH1, we performed lysine-specific cross-linking mass spectrometry (CXMS) using purified βarr1 and SH1. CXMS provides proximity restraints between intermolecular lysine residues and between lysines and proteins N-termini (Fig. 5a, b). Two cross-linked peptides were identified between the N-terminal region of βarr1 and K356 within the C-lobe of the SH1 domain, positioned on the face opposite to the active site (Fig. 5c). To validate the CXMS data, we tested SH1 binding to different βarr1 constructs (full-length, N-domain, and C-domain) by pull-down assay. Whereas the full-length and N-domain of βarr1 bound to SH1, the βarr1 C-domain (residues 176-418) showed no detectable binding to SH1, indicating that the interaction is mediated by the N-domain, in agreement with the cross-linking results (Fig. 5d). The structural data and the proximity restraints from cross-linking analysis suggest that βarr1 engages either SH3 or SH1, but not both simultaneously (Fig. 5e). We propose that inactive βarr1 interacts with Src through SH1; however, upon activation, this interaction appears weakened or lost, and binding shifts in favor of SH3. Further work will be needed to confirm this hypothesis.

**Fig. 5.**
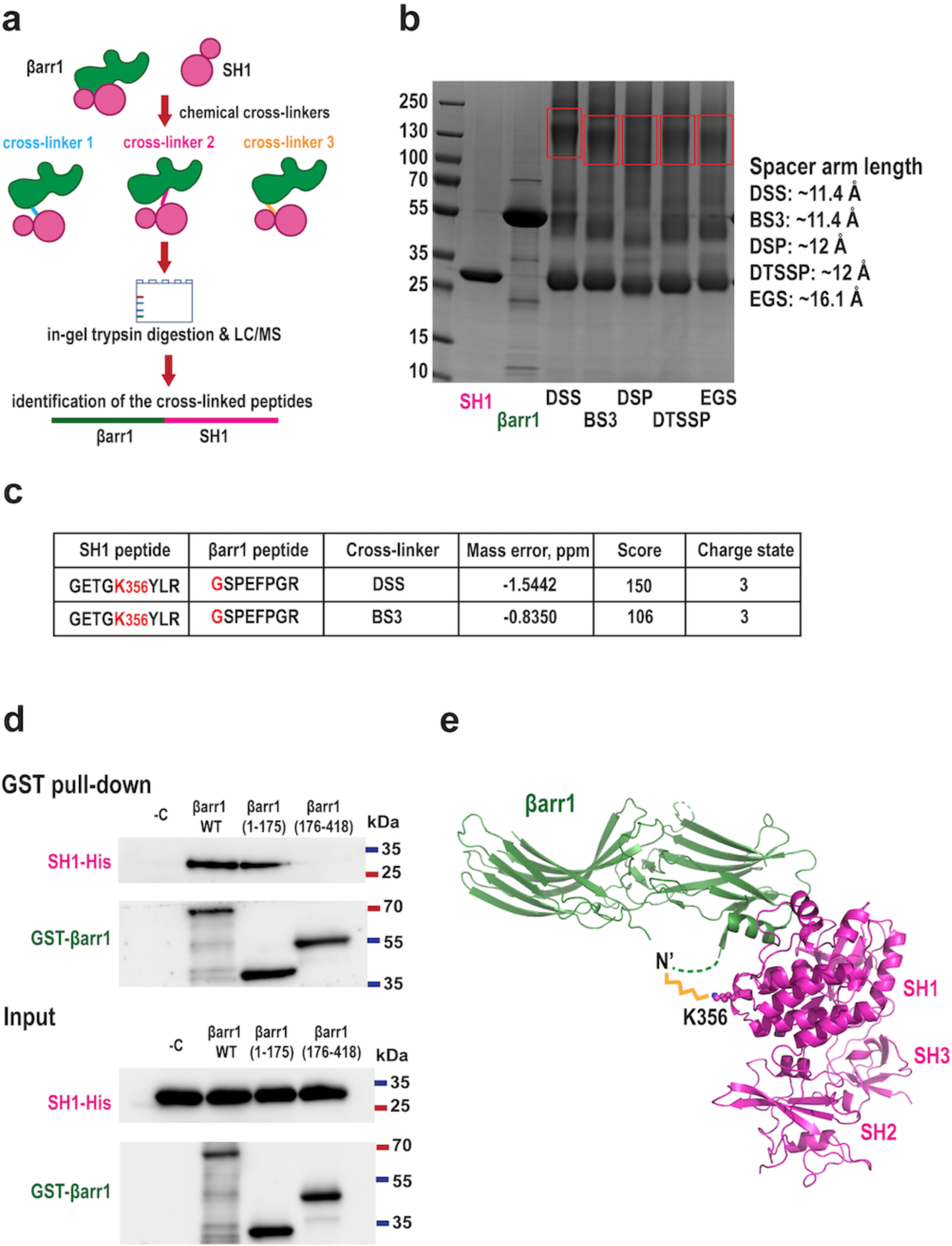
N-domain of βarr1 interacts with SH1. **a,** Schematic of CXMS. **b,** Chemical cross-linking of purified SH1 and βarr1, Coomassie blue gels. The regions of the gels excised for in-gel trypsin digestion are framed in red rectangle. **c**, Identified intramolecular cross-linking peptides (*MaxQuant* score cutoff =100). **d,** Representative Western blot of GST pull-down assay of different constructs of GST-βarr1 (full-length, N-domain (1-175), C-domain (176-418)) and SH1 (n=8). **e,** Model of βarr1 (PDB: 1G4M) binding to the SH1 domain of Src (PDB: 1FMK) based on CXMS data. The cross-link between K356 of SH1 and N-terminus of βarr1 is shown in yellow.

### SH3 binding induces conformational changes in βarr1

Interestingly, βarr1 not only induces conformational changes in Src but also undergoes structural rearrangements upon Src SH3 binding (Fig. 6). The most dramatic changes are observed in the central crest region of βarr1 in the Src–βarr1-CC and SH3–βarr1-CC complexes (Fig. 6a). SH3 binding to βarr1-CC drastically changes the position of β-strand V: it shows a downward and inward movement by ∼9 Å and displays a two-residue offset in its N-terminal part and a three-residue offset in its C-terminal part as compared to active βarr1 without SH3 (Fig. 6a-b). Furthermore, the middle loop in SH3–βarr1-CC adopts an intermediate conformation between the fully active and fully inactive βarr1 crystal structures (Fig. 6c). While we cannot rule out that the differences in the conformation of the middle loop may come from crystal packing constraints or the allosteric effects from binding to GPCRs^24–27^, it is tempting to speculate that the larger conformational changes in β-strand V upon Src SH3 binding might affect the coupling of βarr1 to the receptor and the downstream signaling. We also observed conformational changes in βarr1 within the SH3–βarr1-N complex in proximity to the interaction interface. Specifically, the ^45^PEYLKER^51^ loop shifts by ∼6-8 Å toward SH3 and adopts a distinct conformation compared to that seen in the SH3–βarr1-CC complex and other active βarr1 structures (Fig. 6d). Furthermore, the ^87^FPPAPEDK^94^ motif is shifted by 4 Å toward SH3, although it remains unclear whether this change is due to disulfide cross-linking or SH3 binding (Fig. 6e). Notably, the interaction of βarr1 with Src also introduces local conformational changes in SH3 by affecting the position of the nSrc loop and converting the 3_10_ helix into a loop (Fig. 6f).

**Fig. 6.**
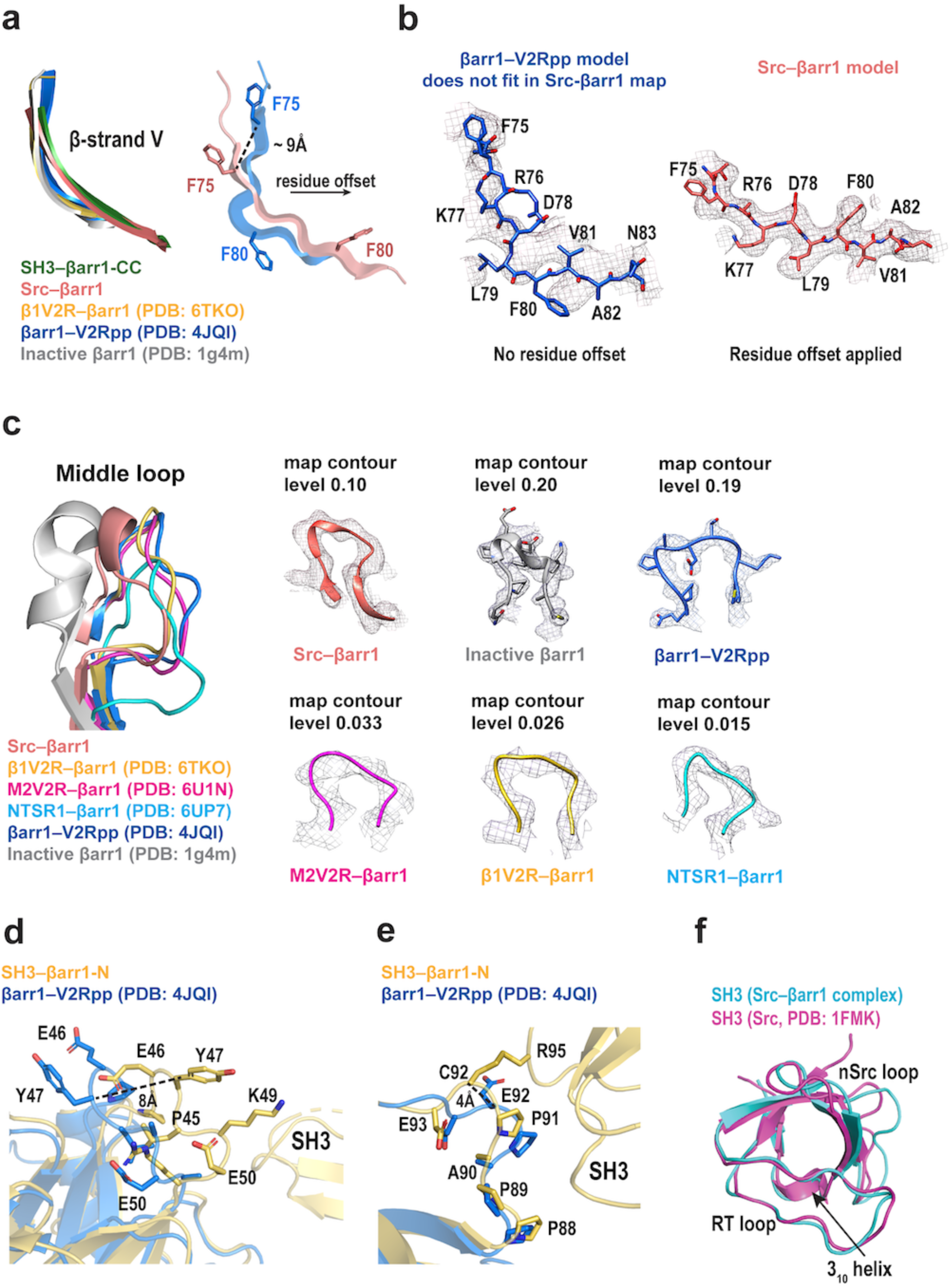
SH3 binding induces distinct conformational changes in βarr1 and SH3. **a-c,** Structural comparison of βarr1 in SH3–βarr1-CC (green), Src–βarr1-CC (salmon), inactive βarr1 (PDB: 1G4M, grey) and active βarr1 structures in complex with: chimeric β1AR with V2R tail (β1V2R) (PDB: 6TKO, yellow), chimeric M2R with V2R tail (M2V2R) (PDB: 6U1N, magenta), neurotensin receptor 1 (NTSR1) (PDB: 6UP7, cyan), V2Rpp–Fab30 (PDB: 4JQI, blue). **a,** Focus on β-strand V. Left panel: superposition of inactive and active βarr1 structures, SH3–βarr1-CC, and Src–βarr1-CC. Right panel: residue offset in Src–βarr1-CC is shown in comparison with βarr1– V2Rpp–Fab30 (PDB: 4JQI, blue); the distance between Cα atoms of F75 is shown. **b,** Left panel: β-strand V of the βarr1–V2Rpp–Fab30 model does not fit into the Src–βarr1-CC density map. Right panel: the residue offset of β-strand V improves the fit into the density. The upsample map is used (0.72 Å/pix); map contour level is 0.1. **c**, Focus on the middle loop. Left panel: superposition of inactive and active βarr1 structures and Src–βarr1-CC. Right panel: density around the middle loop in all respective structures. **d-e,** Structural comparison of βarr1 in SH3– βarr1-N (yellow) and active βarr1 structures in complex with V2Rpp–Fab30 (PDB: 4JQI, blue). Focus on ^45^PEYLKER^51^ loop (**d**) and ^87^FPPAPEDK^94^ motif (**e**). The distances between Cα atoms of Y47 (**d**) and E92/C92 (**e**) are shown. **f,** Structural superposition of SH3 domains in Src–βarr1-CC complex (cyan) and in the crystal structure of Src (PDB: 1FMK, magenta).

### Physiological roles of SH3-binding sites of βarr1

We next sought to elucidate the physiological consequences of SH3 binding to βarr1-N and βarr1-CC. We generated four mutants in the βarr1-N and βarr1-CC sites and tested Src activation in HEK-293 βarr1/βarr2 dKO (CRISPR-Cas9-based βarr1/βarr2 double knock-out) cells^28^. βarr1 mutants showed a decrease in Src activation after stimulation of chimeric β2-adrenergic receptor with V2R tail (β2V2R) with BI-167107 (Fig. 7a). We also tested the activation of endogenous Src in HEK-293 βarr1/βarr2 dKO cells downstream of dopamine 1 (D1R) receptor by βarr1 mutants (Fig. 7b). We chose D1R because Src is one of the key effectors of D1R signaling^29^, and its activation was shown to exclusively depend on βarr^30^. While basal levels of phospho-Src were detected in all non-stimulated cells, addition of dopamine led to a significant increase in phospho-Src in βarr1 WT transfected cells. In contrast, none of the βarr1 mutants showed a response to dopamine stimulation, suggesting their reduced ability to activate Src (Fig. 7b). Therefore, both βarr1-CC and βarr1-N sites mediate Src activation in HEK293 cells downstream of two receptors, β2V2R and D1R. To identify the contribution of individual βarr1 residues on Src activation, we generated six βarr1 mutants and measured the βarr1-mediated activation of Src *in vitro*. Whereas βarr1 WT (activated by V2Rpp and Fab30) increases Src activity two-fold, βarr1-N mutant P88G_P91G, βarr1-CC mutants P121E_P124G_F75A_N122A and P121E_P124G_F75A_I314A, as well as P88G_P91G_P121E_P124G mutant failed to activate Src (Fig. 7c). βarr-CC mutants P121E_P124G and F80A only slightly reduced Src activation compared to βarr1 WT. Therefore, this supports a mechanism where P88 and P91 drive Src activation via the βarr1-N site and F75, N122 and I314 are critical for Src activation via the βarr1-CC site.

**Fig. 7.**
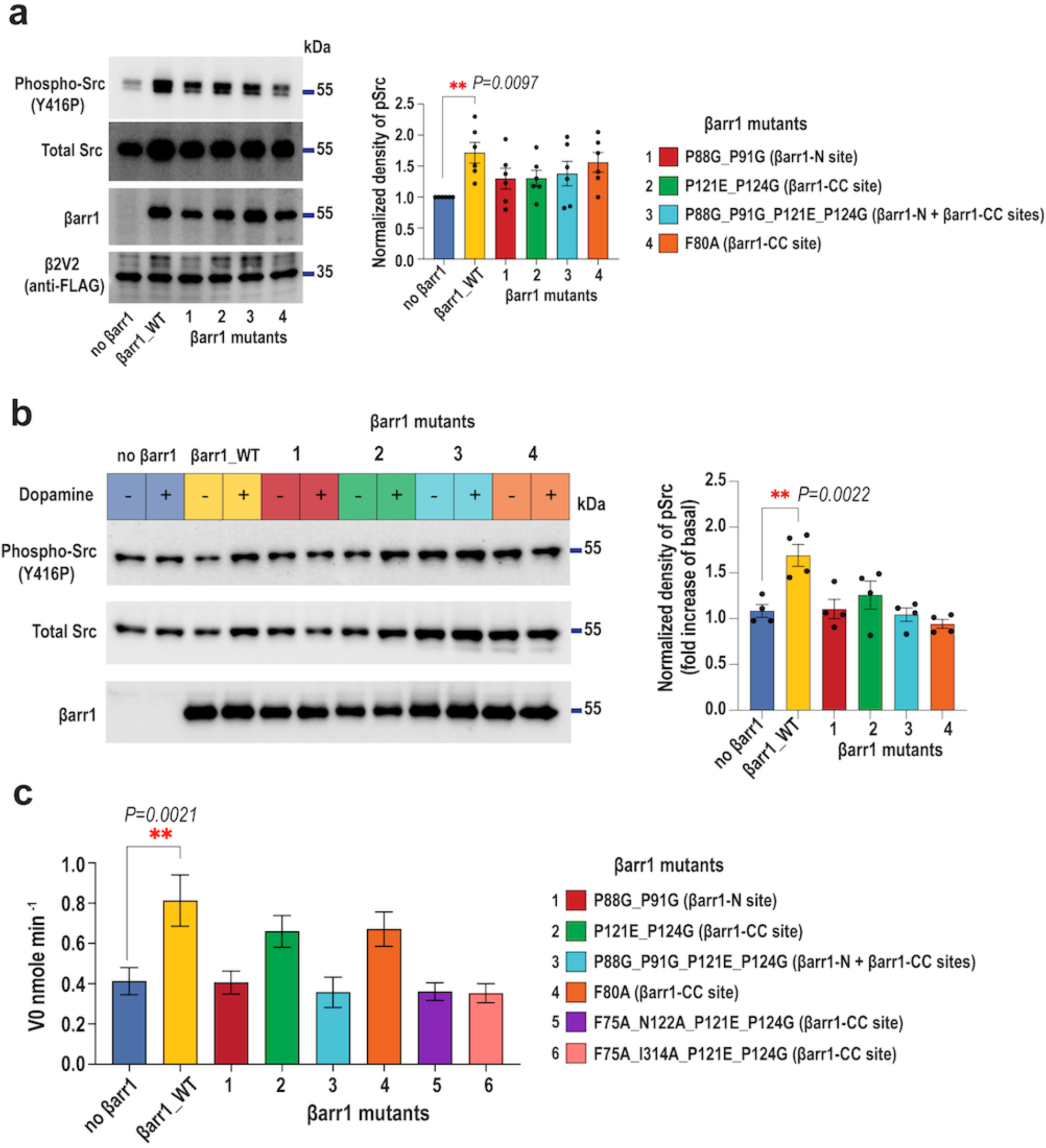
Both βarr1-CC and βarr1-N sites mediate Src activation *in vitro* and in cells. **a,** Effect of βarr1 mutations on Src activity downstream of β2V2. β2V2, Src and βarr1 were transiently expressed in HEK-293 βarr1/βarr2 dKO cells. Representative Western blots and densitometry analysis of Src phosphorylation normalized to phospho-Src in control cells (no βarr1 transfected) (mean ±SEM; n=6; one-way ANOVA with Dunnett’s post hoc test; \*\**P* < 0.01). **b,** Effect of βarr1 mutations on endogenous Src activity downstream of D1R. D1R and βarr1 were transiently expressed in HEK-293 βarr1/βarr2 dKO cells. Representative Western blots and densitometry analysis of Src phosphorylation normalized to phospho-Src in unstimulated cells (mean ±SEM; n=4; one-way ANOVA with Dunnett’s post hoc test; \*\**P* < 0.01). βarr1 mutants numbering is the same as in (**a**). **c,** Initial velocity of optimal Src peptide phosphorylation by Src (V0) alone or in the presence of βarr1 WT or βarr1 mutants (mean ±SEM; n=10; one-way ANOVA with Dunnett’s post hoc test; \*\**P* < 0.01).

We hypothesized that binding of the Src SH3 domain to the central crest region of β-arrestin1 (βarr1-CC), along with the resulting conformational changes in βarr1, could impair its engagement with the transmembrane core of GPCRs (Fig. 8a). We, therefore, evaluated whether excess of SH3 would interfere with the core coupling of βarr1 to the phosphorylated V2R in membranes. We labeled the finger loop of βarr1 V70C with an environment-sensitive bimane fluorophore (mBr) and measured the changes in fluorescence emission spectra with and without SH3 (Fig. 8b). Upon stimulation of the phosphorylated V2R with arginine-vasopressin peptide (AVP), the finger loop of βarr1 V70C-mBr binds to the receptor core, leading to a ∼50% increase in bimane fluorescence. The presence of SH3 significantly reduces this effect, suggesting that SH3 binding to βarr1 sterically hinders the core coupling of βarr1 to V2R. Similar results were obtained with the purified phosphorylated chimeric M2 muscarinic receptor with V2R tail (M2V2R) reconstituted in MSP1D1E3 nanodiscs (Fig. 8c). These data suggest that high local concentration of Src can directly influence the conformational equilibrium of GPCR–βarr1 complexes by disrupting the fully engaged “core” conformation (Fig. 8d), which, in turn, may impair receptor desensitization and hinder the termination of G protein–mediated signaling. To test this hypothesis in a cellular context, we monitored real-time cAMP dynamics in HEK293 cells stably expressing the β2-adrenergic receptor (β2AR) using the live-cell FRET-based ICUE2^31^ biosensor, which consists of a fusion between cyan and yellow fluorescent proteins (CFP-Epac2-YFP) (Fig. 8e). Stimulation of Gαs-coupled β2AR with isoproterenol triggers a rapid peak in cAMP levels, followed by a gradual decline as a result of β2AR desensitization. Overexpression of Src led to increased agonist-induced cAMP accumulation whereas the decline of cAMP accumulation was delayed compared to control cells, indicating that β2AR desensitization is affected (Fig. 8f).

**Fig. 8.**
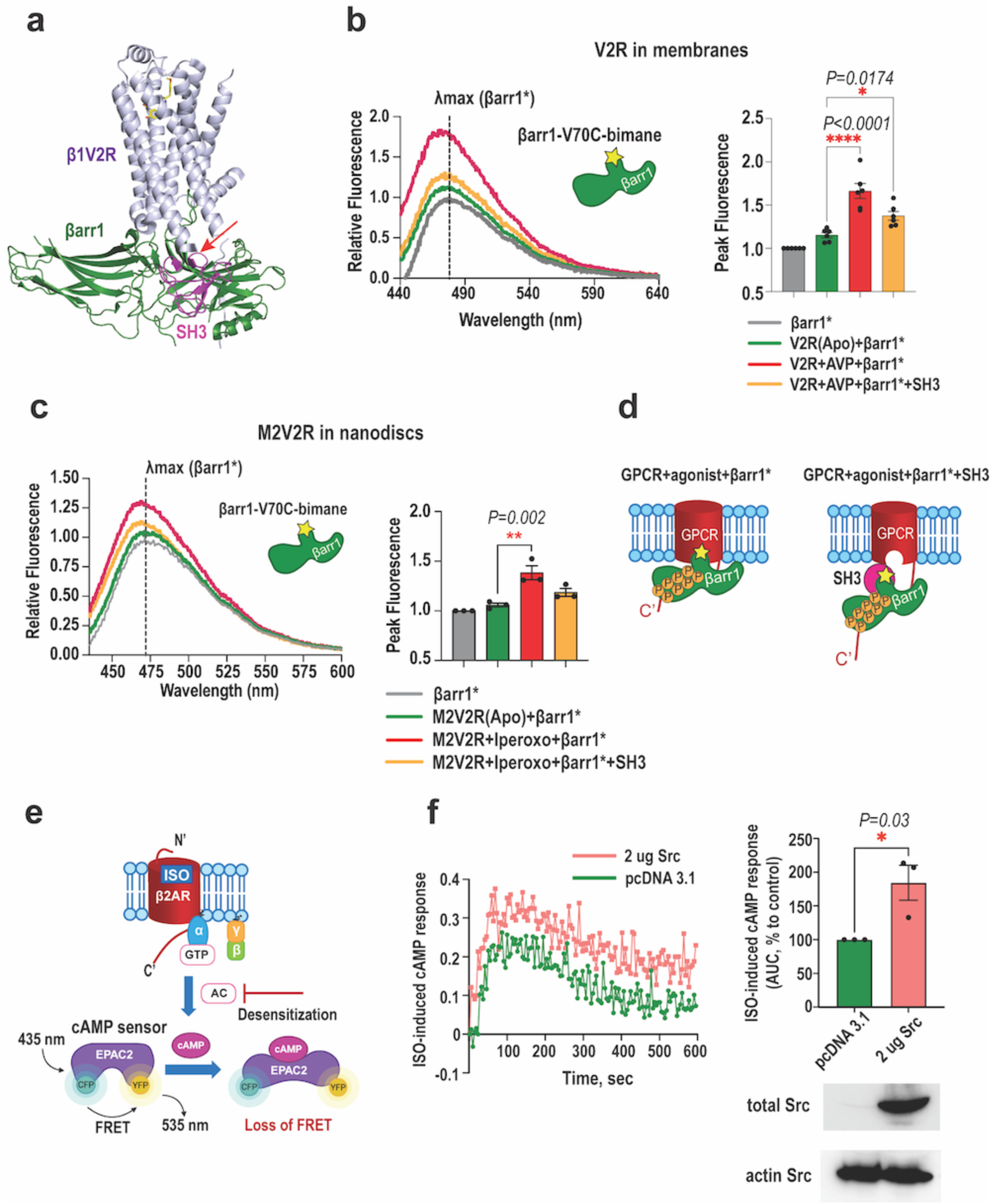
Excess of Src SH3 interferes with the core coupling of βarr1 to GPCRs and overexpression of Src affects desensitization of β2AR. **a,** Structural superposition of Src–βarr1-CC (SH3, magenta; βarr1, green) with the β1V2R–βarr1 complex (β1V2R, slate; PDB: 6TKO), cartoon representation. The clashing region is indicated with the red arrow. **b,** Core coupling of βarr1 finger loop bimane to phosphorylated V2R in membranes with and without SH3. Left panel: representative experiment of βarr1 finger loop bimane fluorescence. Right panel: peak fluorescence of βarr1 finger loop bimane (mean ±SEM; n=6; one-way ANOVA with Dunnett’s post hoc test; \**P* < 0.05, \*\*\*\**P* < 0.0001). **c,** Core coupling of βarr1 finger loop bimane to the phosphorylated chimeric M2V2R reconstituted in MSP1D1E3 nanodiscs. Left panel: representative experiment of βarr1 finger loop bimane fluorescence. Right panel: peak fluorescence of βarr1 finger loop bimane (mean ± SEM; n=3; one-way ANOVA with Dunnett’s post hoc test; \*\**P* < 0.01). **d,** Schematic of SH3-mediated disruption of βarr1 core coupling to GPCRs. **e,** Schematic of the real-time Epac2-FRET cAMP accumulation assay. cAMP binding leads to increased CFP–YPF distance and reduces FRET efficiency. **f**, Left panel: representative kinetic profiles from HEK293 cells stably expressing β2AR and Epac2 and transiently transfected with 2 ug Src or control DNA. Right panel, top: area under the curve (AUC) of FRET responses relative to control (100%). Right panel, bottom: representative Western blots of cell lysates.

## Discussion

GPCRs convert extracellular signals into cellular responses via two major transducers, G proteins and βarrs, which in turn interact with their downstream effectors. The rapid development of cryo-EM has greatly advanced our understanding of how GPCRs recruit signal transducers. In contrast, little is known about how the transducers regulate their effector enzymes. Protein interactions within signaling cascades are inherently weak and transient, which is crucial for the dynamic regulation of cellular processes. Capturing dynamic and transient protein complexes has long posed a significant challenge in structural biology, often requiring stabilization strategies such as genetic fusions, chemical cross-linking, disulfide trapping, or extensive mutagenesis to obtain structural information that would otherwise be unattainable^32–34^. Among these methods, disulfide trapping offers distinct advantages: it minimizes non-specific artifacts compared to chemical cross-linking and has a lower risk of disrupting protein folding or expression compared to genetic fusion approaches. Using disulfide trapping, we determined the first high-resolution structures of a βarr-effector signaling hub, βarr1 in complex with tyrosine kinase Src and isolated SH3 bound to two distinct sites of βarr1. The structural data reveal a unique strategy utilized by βarr1 to recruit Src SH3. βarr1 binds the aromatic surface of SH3 using two distinct sites, each with its own binding mechanism. The βarr1-N site is a short polyproline motif, whereas the βarr1-CC site is formed by the non-proline residues in the β-strand V and the lariat loop and two proline residues.

The unique ability of βarr1 to engage its effector proteins through two distinct sites using different binding mechanisms highlights a sophisticated level of modularity and flexibility in signaling scaffolds. This dual-site architecture likely plays a central role in the remarkable functional versatility of βarr1 across diverse cellular contexts. Several key implications emerge from this mode of engagement. First, this configuration enables βarr1 to ensure robust recruitment and activation of Src when one of the interaction sites is sterically hindered or occupied by a competing binding partner. Second, two SH3-binding surfaces could also facilitate the simultaneous or sequential recruitment of effectors, enabling the assembly of multi-component signaling complexes or temporally orchestrated signaling cascades. In the context of Src, βarr-mediated conformational activation at two distinct sites may also promote Src trans-autophosphorylation, as the two kinase molecules are brought into close proximity. In proteins containing multiple SH3 domains, dual-site engagement can enhance the overall avidity of the interaction, stabilizing the complex. Third, the dual-site architecture might contribute to the spatiotemporal compartmentalization of Src signaling^35^, with one site functioning predominantly at the plasma membrane and the other in endosomal compartments. Such spatial regulation could underlie distinct modes of Src activation (transient vs. sustained) and lead to different signaling outcomes, such as rapid cytoskeletal rearrangement or longer-term transcriptional responses. Finally, different upstream signals may induce conformational states of βarr1 that preferentially expose one Src-binding site over the other. While our data show that signaling downstream of β2V2 and D1R did not display a clear bias toward either site, this may not be the case for other GPCRs or in response to different ligands, presenting an exciting direction for future research. Furthermore, post-translational modifications of βarr1 could influence the accessibility or affinity of each site, allowing βarr1 to function as a dynamic molecular gate with context-dependent control over Src activation. Beyond Src, this dual-recognition strategy may represent a broader mechanism by which βarr1 engages other SH3 domain-containing proteins. Elucidating the biological consequences of the dual-site recognition mechanisms of βarr1 opens exciting new avenues for exploring its complex regulatory roles in cellular signaling.

Src activation by SH3/SH2-binding proteins is well established: binding to these domains disrupts the intramolecular interactions that constrain the kinase activity^36–42^. Prior studies suggested that βarr1 activates Src via SH3 domain interaction^3, 4^. Using site-specific fluorescence labeling of Src, Yang *et al.* showed the disruption of SH3/SH1 and SH3/SH2 interactions in the presence of β2AR-βarr1 complex^3^. However, in the absence of structural data, it remained unclear whether these effects result from βarr1’s scaffolding or its direct activation of Src. Here, we present the first structural evidence that βarr1 functions as an active regulatory protein rather than a passive scaffold and elucidate the precise molecular mechanism of βarr1-mediated allosteric activation of Src (Fig. 4, Supplementary Video 3). At both sites, βarr1 interacts with the aromatic surface of SH3, which is critical for maintaining the autoinhibited conformation of Src. This interaction disrupts the autoinhibitory interactions, liberating the catalytic domain of Src, as observed in the structure of the Src–βarr1-CC complex.

Beyond allosteric activation, βarrs might also facilitate trans-autophosphorylation of Src by bringing two Src molecules into close proximity. Furthermore, βarrs may position Src near its substrates, analogous to the arrangement seen in focal adhesion kinase (FAK)–Src signaling complexes^43^.

Notably, Src SH3 and βarr1 exhibit reciprocal effects: βarr1 induces conformational changes in Src promoting its activation, while Src SH3 simultaneously triggers significant structural rearrangements in βarr1. The conformational changes in the central crest region of βarr1 upon SH3/Src binding may affect βarr1 interactions with GPCRs and other signaling partners (Fig. 6). Our results demonstrate that an excess of SH3 interferes with βarr1 coupling to the receptor core (Fig. 8b-d). Furthermore, in HEK293 cells overexpressing Src, we observed elevated agonist-induced cAMP accumulation, indicating impaired β2AR desensitization (Fig. 8e, f). Although the βarr1–Src interaction is likely transient, a high local concentration of Src may shift the conformational equilibrium of GPCR–βarr1 complexes toward a “tail-only” engagement, which permits sustained G-protein signaling. This mechanism is reminiscent of sustained G-protein signaling from internalized GPCRs via megaplex assemblies, in which GPCRs simultaneously engage both G-proteins through its core region and βarr1 through its phosphorylated C-terminal tail^44, 45^. Future structural studies on the GPCR–βarr–Src axis will be crucial to define the conformational states of the complex and assess their physiological relevance. A promising direction is to test the implications of this mechanism in cancer models, where Src is frequently overexpressed^46^, to understand its role in disease progression and signaling dysregulation. Our results suggest that Src overexpression may disrupt receptor desensitization.

βarrs, originally discovered as proteins mediating GPCR desensitization^47, 48^, are now commonly recognized as signal transducers^2–4, 30^. Nevertheless, some controversy remains as to whether βarrs are just modulators of G-protein-mediated signaling events^49^ or rather *bona fide* signal transducers in their own right. Our cryo-EM structures support the latter: βarr1 recruits Src and allosterically activates the enzyme by disrupting its autoinhibitory intramolecular interactions. This may represent a general mechanism by which βarrs mediate signal transduction across diverse effectors. Taken together, our study demonstrates how βarrs orchestrate cellular signaling downstream of GPCRs and highlights new avenues for exploring their regulatory roles in both physiological and pathological contexts.

## Methods

### Bacterial strains

*Escherichia coli* strain DH5α (C2987, New England Biolabs) was used for plasmid propagation. Protein expression was done in *E. coli* BL21 derivative strain (C3010, New England Biolabs).

### Cell culture

Human embryonic kidney CRISPR-Cas9-based βarr1/βarr2 double knock-out cell line (HEK-293 βarr1/βarr2 dKO)^28^ was maintained in Gibco Minimum Essential Media (MEM) supplemented with 1% penicillin-streptomycin and 10% (v/v) fetal bovine serum at 37 °C and 5% CO_2_.

### Molecular biology

Bacterial expression constructs for wild-type (residues 2-418) and a minimal cysteine (C59A, C125S, C140I, C150V, C242V, C251V, and C269S) and truncated (βarr1–MC-393) variants of rat βarr1,^50^ Fab30,^51^ and Nb32^11^ have been reported previously. Plasmid pET28a-3D-Src expressing SH3, SH2 and SH1 domains of chicken Src (residues 83-533), plasmids expressing YopH phosphatase and Csk were generous gifts from Prof. John Kuriyan. For structural studies Src-5Cys mutant (C185S, C238S, C245S, C277S, C400S) was generated. Bacterial expression constructs for Src SH1 domain (250-536) and for βarr1 C-domain (177-420) were synthesized by GenScript. Plasmid expressing the SH3 domain of Src (residues 83-141) was designed by inserting a C-terminal FLAG-tag and a stop codon in pET28a-3D-Src. Mammalian expression constructs for human FLAG-M2-muscarinic receptor (M2R) with C-terminal sortase ligation consensus sequence (LPETGGH) and 6×His-tag,^50^ chimeric FLAG-β2-adrenergic receptor with C-terminal tail of vasopressin 2 receptor (β2V2),^52^ SNAP-vasopressin 2 receptor (V2R),^44^ HA-tagged βarr1,^9^ and human Src^9^ have been previously reported and functionally verified. Mammalian expression construct for HA-dopamine 1 receptor (D1R) was a gift from Prof. Marc Caron. Mutations were introduced using QuickChange II site-directed mutagenesis kit (Agilent) and verified by Sanger sequencing.

### Protein expression and purification

The expression and purification of Src from *Escherichia coli* are described in detail elsewhere ^53^. Src construct (residues 83-533) without the N-terminal disordered region was used for the structural and *in vitro* studies. Briefly, Src was co-expressed with YopH phosphatase and purified by immobilized metal ion affinity chromatography and anion exchange chromatography. The SH3 domain of Src and its variants were purified by immobilized metal ion affinity chromatography and size exclusion chromatography (SEC). Wild-type βarr1 and its variants,^54^ Fab30,^55^ and Nb32^11^ were expressed and purified as described previously. The expression and purification of FLAG-M2 receptor containing a C-terminal sortase ligation consensus sequence (LPETGGH), sortase ligation reaction and reconstitution in MSP1D1E3 nanodiscs are described elsewhere.^25, 50^

### Disulfide trapping

βarr1–MC-393 mutant (30 μM) was incubated with 3-fold molar excess of V2Rpp, 1.5 molar excess of Fab30, and 2-fold molar excess of SH3 mutant or Src mutant in 20 mM HEPES 7.5, 150 mM NaCl buffer for 30 minutes at room temperature. The disulfide trapping reactions were initiated by 1 mM H_2_O_2_. After 1-hour incubation at room temperature the reactions were terminated by addition of 4x Laemmli sample buffer (without β-mercaptoethanol) (BioRad), subjected to SDS-PAGE, visualized by Ready Blue Coomassie stain (Sigma-Aldrich) and quantified by *ImageJ v1.52a*. The band of SH3–βarr1 complex (∼60 kDa) or Src–βarr1 (∼130 kDa) was normalized to the total density of all bands in each sample. To test the effect of V2Rpp and Fab30 on formation of disulfide trapped complexes, the reactions were performed as described above; in reactions without V2Rpp and Fab30 the equivalent amount of buffer was added. The samples were subjected to SDS-PAGE and Western blotting, detected by polyclonal A1CT antibody generated in Lefkowitz lab^56^ (1:5000) for βarr1.

### Isothermal titration calorimetry (ITC)

ITC measurements were performed using the MicroCal PEAQ-ITC system (Malvern). Purified βarr1–V2Rpp and SH3 were dialyzed in 20 mM HEPES, pH 7.5, 100 mM NaCl. 75 μM of βarr1– V2Rpp was loaded into the sample cell and 750 μM of SH3 into the injection syringe. The system was equilibrated to 25 °C. Titration curves were initiated by a 0.4 μL injection from syringe, followed by 2.0 μL injections (at 180 s intervals) into the sample cell. During the experiment, the reference power was set to 7 μcal·s-1 and the sample cell was stirred continuously at 750 rpm. Raw data, excluding the peak from the first injection, were baseline corrected, peak area integrated, and normalized. Data were analyzed using *MicroCal Origin* software to obtain thermodynamic parameters of binding and association constant (Ka=1/Kd).

### Formation of SH3–βarr1 and Src–βarr1 complexes

The βarr1–MC-393 mutant (100 μM) was incubated with 2-fold molar excess of the SH3 mutant or Src mutant for 30 minutes at room temperature, followed by disulfide trapping as described above. SH3–βarr1 was separated from unbound βarr1 by immobilized metal ion affinity chromatography using Talon resin (Takara Bio). Src–βarr1 was separated from unbound βarr1 and Src by SEC. SH3–βarr1 and Src–βarr1 complexes were then incubated with 3-fold molar excess of V2Rpp, 1.5-fold molar excess of Fab30, 2-fold molar excess of Nb32 (for SH3–βarr1-CC and Src–βarr1-CC) for 30 minutes at room temperature. The complexes were subjected to SEC on a Superdex 200 Increase column (Cytiva Life Sciences) in 20 mM HEPES 7.5, 150 mM NaCl buffer. Peak fractions were concentrated to 6-8 mg/ml using Vivaspin® 6 column with molecular weight cut-off of 30,000 kDa (Sartorius).

### Cryo-EM grid preparation and data acquisition

In the preliminary cryo-EM experiments, we observed that the complexes exhibited strong preferred orientation in vitreous ice. To alleviate the preferred orientation problem, 8 mM CHAPSO was added to the sample immediately before vitrification.^57^ The sample was applied to glow-discharged 300-mesh holey-carbon grids (Quantifoil R1.2/1.3, Electron Microscopy Sciences) or gold grids (Ultrafoil R0.6/1.0, Electron Microscopy Sciences) and vitrified using a Vitrobot Mark IV (Thermo Fisher Scientific) at 4 °C and 100% humidity. The data were collected on a Titan Krios transmission electron microscope (Thermo Fisher) operating at 300 kV equipped with a K3 direct electron detector (Gatan) in counting mode with a BioQuantum GIF energy filter (slit width of 20 eV) at a magnification of ×81,000 corresponding to a pixel size of 1.08 Å at the specimen level. 60-frame Videos with a dose rate of ∼15 electrons per pixel per second and a total accumulated dose of ∼53-60 electrons per Å^2^ were collected using the *Latitude-S* (Gatan) single-particle data acquisition program. The nominal defocus values were set from −0.8 to −2.5 µm.

### Cryo-EM data processing

Videos were subjected to beam-induced motion correction using Patch Motion Correction in *CryoSPARC v4.0.1*^58^ followed by determination of Contrast transfer function (CTF) parameters in Patch CTF. Micrographs with CTF fit better than 3.5 Å were used for further analysis. Particles manually selected from 15 micrographs were used to train a model in the particle picking tool *Topaz v0.2.5a*.^59^ The trained model was used to pick particles in all micrographs generating 200,270 particle projections for the SH3–βarr1-CC complex and 760,233 particle projections for the SH3–βarr1-N complex. The particles were rescaled to the pixel size of 1.3824 Å for further processing. For the SH3–βarr1-CC complex, the particle stack was subjected to 2D classification, *Ab Initio* model generation, non-uniform refinement and local refinement with a mask excluding the constant (CL/CH1) region of Fab30 (FabCR) in *CryoSPARC*. Particles were then imported to *RELION v3.1*^60^ and subjected to 3D classification without alignment with a mask excluding the constant (CL/CH1) region of Fab30 (FabCR). Classes showing a clearly defined density for the SH3 domain (118,020 particles) were re-imported to *CryoSPARC* and subjected to local refinement with a mask on FabCR and a fulcrum on the SH3–βarr1 interface (Supplementary Fig. 3a). The resulting map was then processed by spIsoNet to improve map isotropy^61^. The final map has a global resolution of 3.47 Å at a Fourier shell correlation of 0.143 (Supplementary Fig. 4c). For the SH3–βarr1-N complex, the particles picked with *Topaz* were subjected to 2D classification and the best 2D classes were used in template picking (5,607,258 particles). A subset of particles (1,000,000) was used to generate six *Ab Initio* classes. All particles were then subjected to four rounds of heterogeneous refinement. The resulting particle stack (345,529) was used in non-uniform refinement and local refinement with a mask excluding FabCR (Supplementary Fig. 3b). The resulting map was then subjected to spIsoNet to improve map isotropy^61^. The final map has a global resolution of 3.34 Å at a Fourier shell correlation of 0.143 (Supplementary Fig. 4d). For the Src–βarr1-CC complex, the particles picked with *Topaz* (94,196 particles) were subjected to 2D classification and the best 2D classes were used in template picking (9,770,378 particles). The particles were rescaled to the pixel size of 1.44 Å for further processing. A subset of particles (1,000,000) was used to generate three *Ab Initio* classes. All particles were then subjected to three rounds of heterogeneous refinement, followed by non-uniform refinement and local refinement with a mask excluding FabCR. The resulting particle stack (692,850) was subjected to 3D classification without alignment in *CryoSPARC* and then in *RELION* with a mask excluding FabCR. Finally, the best class with 140,156 particles was subjected to another round of local refinement in *CryoSPARC,* generating a map with a global resolution of 3.34 Å (Supplementary Fig. 7c, Supplementary Fig. 8b). For the Src–βarr1-N complex, the particles picked with *Topaz* (61,962 particles) were subjected to 2D classification and matched with the 2D projections of the SH3–βarr1-N map using cluster selection mode in Reference Based Auto Select 2D module in *CryoSPARC*.

### Model building and refinement

Cryo-EM maps for SH3–βarr1 complexes were post-processed using DeepEMhancer^62^ to improve the interpretability of maps, specifically in the βarr1 loops and at the SH3–βarr1 interface. Since the overall resolution of maps was better than 4 Å, highRes deep learning model was used during DeepEMhancer processing of maps. The initial models of the SH3–βarr1-CC complex and Src– βarr1-CC complex were built manually by fitting the crystal structures of SH3 (PDB: 2PTK), βarr1–V2Rpp–Fab30–Nb32 complex (PDB code: 6NI2) into the experimental electron densities using *UCSF Chimera v1.15.*^63^ The structures were refined by combining manual adjustments in *Coot v0.9.8.3*^64^ and ISOLDE^65^ in *UCSF ChimeraX 1.6.1,*^66^ followed by real-space refinement in *PHENIX v1.20.1-4487*^67^ with Ramachandran, rotamer, torsion, and secondary structure restraints enforced. For the SH3–βarr1-N complex, the crystal structure of the βarr1–V2Rpp–Fab30 complex (PDB code: 4JQI) was first built manually into the density and refined similarly to the SH3–βarr1-CC complex. After the refinement, SH3 (PDB: 2PTK) was docked into the density guided by the position of the disulfide bond between βarr1_E92C and SH3_R95C. The rigid body fitted SH3– βarr1-N model generated above was then subjected to molecular dynamics flexible fitting (MDFF) using Cryo fit,^68^ integrated in Phenix software suite and Namdinator^69^ followed by further iterative rounds of model building and real space refinement in *Coot v0.9.8.3*^64^ and *PHENIX v1.20.1-4487.*^67^ Due to weak density for the SH3 domain in the SH3–βarr1-N complex, only its main chain was modeled. The models were validated with *MolProbity v4.5.1*.^70^ Refinement statistics are given in Supplementary Table 1.

### Hydrogen-deuterium exchange mass-spectrometry

To analyze the solvent accessibility of βarr1, βarr1–MC-393 (20 μM) or βarr1–MC-393–V2Rpp (20 uM βarr1; 100 uM V2Rpp) were incubated with 5-fold molar excess of SH3 in the 20 mM HEPES 7.5, 100 mM NaCl buffer for 30 minutes on ice. To test the solvent accessibility of SH3, SH3 (20 uM) was incubated with 5-fold molar excess of βarr1–MC-393 or βarr1–MC-393–V2Rpp in the 20 mM HEPES 7.5, 100 mM NaCl buffer for 30 minutes on ice. The protein stock solution was diluted 17-fold in the 20 mM HEPES 7.5, 100 mM NaCl buffer prepared in D_2_O. At a designated time point (300, 1000, and 5000 s), the exchange was quenched by adding an equal volume of 0.8% formic acid in H2O, yielding pH 2.6. The sample was digested on Enzymate BEH Pepsin column (Waters) at a flow rate of 0.15 ml/min using 0.15% formic acid/3% acetonitrile as the mobile phase. An inline 4-μl C8-Opti-lynx II trap cartridge (Optimize Technologies) was used for desalting of the digested peptides. The peptides were then eluted through a C-18 column (Thermo Fisher Scientific, 50 × 1 mm Hypersil Gold C-18) using a rapid gradient from 10 to 90% acetonitrile containing 0.15% formic acid and a flow rate of 0.04 ml/min, leading directly into a maXis-II ETD ESI-QqTOF mass spectrometer (Bruker Daltonics). The total time for the digest and desalting was 3 min, and all peptides had eluted from the C-18 column by 15 min. Pepsin and C-18 column were thoroughly washed after each run. The peptide fragments were identified using *Bruker Compass* and *Biotools* software packages. The level of deuterium incorporation was calculated using *HDExaminer-3* (Trajan Scientific) from triplicate measurements of each time point. Only regions that exhibited statistically significant differences in deuterium uptake (>0.2 Da) across at least two peptides or two time points were considered relevant.

### M2-FLAG pull-down assay

To test the binding between βarr1 and the SH3 domain of Src, βarr1 (20 μM) was incubated with 3-fold molar excess of V2Rpp for 30 minutes at room temperature, then 10 μM of SH3-FLAG was added. 30 μl of anti-FLAG M2 affinity gel (Millipore Sigma) was added thereafter and the mixture was incubated for 1 hour at room temperature with rotation. After incubation the anti-FLAG M2 resin was collected by centrifugation and washed with 1 ml of 20 mM HEPES pH 7.5, 100 mM NaCl buffer three times. The proteins were eluted with 0.2 mg/ml FLAG-peptide in 20 mM HEPES pH 7.5, 100 mM NaCl buffer, then mixed with Laemmli sample buffer (BioRad), subjected to SDS-PAGE and Western blotting and detected by monoclonal ANTI-FLAG M2-peroxidase (HRP) antibody (1:2000) (A8592, Sigma-Aldrich, RRID: AB_439702) for SH3-FLAG and polyclonal A1CT antibody^56^ (1:5000) for βarr1.

### *In vitro* Src C-tail phosphorylation followed by M2-FLAG pull-down assay

Purified Src (20 μM) was incubated with 4 μM Csk in the presence of 100 μM ATP and 5 mM MgCl2 for 2 hours at room temperature. A control reaction was prepared identically, excluding ATP and MgCl2. The C-tail phosphorylation of Src was assessed by Western blotting using Anti-Src (phospho Y529) antibody (ab32078, Abcam, RRID: AB2286707). The samples of C-tail phosphorylated and unphosphorylated Src (20 μM) were added to 7 μM of βarr1-FLAG pre-incubated with 35 μM of V2Rpp. 30 μl of anti-FLAG M2 affinity gel (Millipore Sigma) was added thereafter and the mixture was incubated for 1 hour at room temperature with rotation. After incubation the anti-FLAG M2 resin was collected by centrifugation and washed with 1 ml of 20 mM HEPES pH 7.5, 100 mM NaCl buffer three times. The proteins were eluted with 0.2 mg/ml FLAG-peptide in 20 mM HEPES pH 7.5, 100 mM NaCl buffer, then mixed with Laemmli sample buffer (BioRad), subjected to SDS-PAGE and Western blotting. The total Src was detected by anti-Src antibody (EMD Millipore 05-184, RRID: AB_2302631); βarr1-FLAG was detected by monoclonal ANTI-FLAG M2-peroxidase (HRP) antibody (1:2000) (A8592, Sigma-Aldrich, RRID: AB_439702). Western Blot images were taken using BioRad ChemiDoc system and the densitometry analysis was performed by *ImageLab v6.1* and statistical differences were determined by Mann-Whitney test in GraphPad Prism software.

### Cross-linking mass-spectrometry

Purified βarr1 (90 μM) was incubated the purified SH1 domain of Src (90 μM) for 1 hour at room temperature. Cross-linking reagents (disuccinimidyl suberate (DSS), Bis(sulfosuccinimidyl) suberate (BS3), dithiobis[succinimidylpropionate] (DSP), 3,3’- dithiobis[sulfosuccinimidylpropionate] (DTSSP), ethylene glycol bis(succinimidyl succinate)) (EGS)) were added thereafter to a final concentration of 2 mM. The reactions were incubated for 30 min and quenched by 50 mM Tris pH 8.0 for 15 min. The proteins were subjected to SDS-PAGE and visualized by Ready Blue Coomassie stain (Sigma-Aldrich). The bands of interest (80-150 kDa) were excised from the gel and subjected to destaining using 25 mM ammonium bicarbonate in 50% acetonitrile. Destained gel slices were dehydrated using 100% acetonitrile and dried using speed-vac for 5 min to remove remaining acetonitrile. 10mM dithiothreitol and 55 mM iodoacetamide were used for reduction and alkylation of cysteine residues, respectively. The gel slices were washed again in 25 mM ammonium bicarbonate and dried using 100% acetonitrile following by 5-min speed-vac evaporation. The proteins were then subject to in-gel trypsin digestion with 10ng/µl trypsin (Sequencing grade, Promega) in 25 mM ammonium bicarbonate and overnight at 37° C. The following day, the tryptic peptides were extracted with 100% acetonitrile and lyophilized. The peptides were loaded onto EvoSep tips (EvoSep) following manufacturer’s instructions. The peptide samples were analyzed using a timsTOF Pro 2 mass spectrometer (Bruker) coupled with an Evosep One LC system (Evosep) connected to an 8-cm Evosep performance column (EV-1109) using a pre-defined 30-samples-per-day extended gradient. The following parameters were applied for data-dependent acquisition: intensity threshold: 2.5E3; charge states: +1 to +5; dynamic exclusion: 0.4 min; target intensity for fragmentation: 2E4; isolation window (linear): 2 m/z at 700 m/z and 3 m/z at 800 m/z; collision energy (linear): 20 eV at 1/k0 of 0.60 V·s/cm2 and 59 eV at 1/k0 of 1.60 V·s/cm2. The cross-linked peptides were identified using *MaxQuant* with a score threshold of 100.

### GST pull down assay

To test the binding between different constructs of GST-βarr1 (full-length βarr1, βarr1-N domain (1-176), βarr1-C domain (177-420)) and the SH1 domain of Src, GST-βarr1 (3 μM) was incubated with SH1 (9 μM) for 1 hour at room temperature. 50 μL of GST-beads (GoldBio) equilibrated in 20 mM HEPES pH 7.5, 100 mM NaCl buffer were added thereafter, and the mixture was incubated for 1 hour at room temperature with rotation. After incubation the GST beads were collected by centrifugation and washed with 20 mM HEPES pH 7.5, 150 mM NaCl buffer three times. The proteins were eluted from GST beads with 100 μL of 50 mM reduced glutathione in 20 mM HEPES pH 8.0, 150 mM NaCl buffer, then mixed with 4x Laemmli sample buffer (BioRad), subjected to SDS-PAGE and Western blotting. SH1 was detected by polyclonal HRP-conjugated Anti-6X His-tag antibody (ab1187, Abcam, RRID: AB_298652); βarr1 was detected by monoclonal HRP-conjugated GST-tag antibody (8-326) (MA4-004-HRP, ThermoFisher, RRID: AB_2537634).

### *In vitro* Src continuous colorimetric kinase assay

Continuous colorimetric kinase assay was performed as previously described.^4, 71^ The reactions were performed in 100 mM HEPES 7.5, 150 mM NaCl, 5 mM MgCl2, 0.005% Triton X-100, containing 0.25 mM optimal Src peptide (AEEEIYGEFEAKKKK), 1 mM phosphoenolpyruvate, 0.3 mM NADH, 4 units of pyruvate kinase and 6 units of lactic dehydrogenase. The concentration of Src was 20 nM, the concentrations of βarr1 and V2Rpp were 100 nM and 200 nM, respectively. Reactions were started by the addition of ATP to a final concentration of 0.1 mM, and the decrease in NADH absorbance was monitored over 40 min at 25° C using a CLARIOstar microplate reader (BMG Labtech). The initial velocity of the reaction (V0) determined using a linear regression curve fit (*GraphPad Prism v9*) and was converted to the amount of product formed in the reaction volume per minute using the Beer-Lambert law. Statistical comparisons were determined by one-way ANOVA with Dunnett’s post hoc test.

### Src activation assay in HEK-293 βarr1/βarr2 dKO cells

HEK-293 βarr1/βarr2 dKO cells^28^ were co-transfected with receptor (FLAG-D1R or chimeric FLAG-β2V2), Src (only for chimeric FLAG-β2V2) and wild-type or mutant βarr1 with a C-terminal HA tag with a 1:5 DNA:FuGENE®6 (Promega) ratio according to the manufacturer’s instructions. All experiments were conducted 48 hours after transfection. Cells were serum starved for 16 hours in MEM supplemented with 1% penicillin-streptomycin and 0.1% (w/v) bovine serum albumin. The assay was initiated with 5-minute stimulation (10 μM dopamine for D1R) or 10-minute stimulation (10 μM BI-167107 for β2V2) at 37 °C. The medium was removed, cells were placed on ice and lysed with 2x Laemmli sample buffer (BioRad) supplemented with 4% β-mercaptoethanol. Samples of equal volume from cell lysates were separated with SDS-PAGE, transferred to nitrocellulose membranes and immunoblotted. Primary antibodies and dilution used are as follow: monoclonal ANTI-FLAG M2 peroxidase (HRP) antibody (1:2000) (A8592, Sigma-Aldrich, RRID: AB_439702) to detect β2V2, polyclonal A1CT antibody generated in Lefkowitz lab^56^ (1:5000) for wild-type and mutant βarr1, anti-Src monoclonal antibody (1:1000) (MA5-15214, Thermo Fisher Scientific, RRID: AB_10980540) for total Src, anti-Src polyclonal Y418 for overexpressed Src (1:5000) (ab4816, Abcam, RRID: AB_304652), and Phospho-Src family polyclonal Y416 for endogenous Src (1:10000) (2101, Cell Signaling, RRID: AB_331697). Secondary antibodies included HRP conjugated anti-rabbit (NA934, Cytiva Life Sciences, RRID: AB_772206) and anti-mouse (NA931, Cytiva Life Sciences, RRID: AB_772210). Western Blot images were taken using BioRad ChemiDoc system and the densitometry analysis was performed by *ImageJ v1.52a* and *ImageLab v6.1*.

### Bimane fluorescence

Purified βarr1-MC-393 V70C was labeled with monobromobimane (mBr) (Sigma-Aldrich), as described previously.^25^ The experiments were performed using membranes with phosphorylated V2R and purified chimeric M2V2R reconstituted in lipid nanodiscs. Expi-293 cells grown in suspension were co-transfected with plasmids expressing the human V2R and the membrane targeted form of GRK2. Transfected cells were grown in suspension for 48h and thereafter were stimulated with AVP (100 nM) for 30 min at 37°C followed by harvesting by centrifugation at 4°C. Cell pellets were washed with cold HBSS, snap frozen in liquid N2 and stored at -80°C prior to preparation of crude membranes as described previously ^72^. Aliquots of phosphorylated V2R membranes were activated with AVP (10μM) then incubated with βarr1 V70C-mBr (20 nM), and SH3 (250 nM) in the assay buffer (20 mM HEPES 7.4, 100 mM NaCl, 1 mg/ml bovine serum albumin) for 120 min at room temperature in black solid-bottom 96-well microplates (Corning) with gentle agitation. Purified M2V2R reconstituted in lipid nanodiscs (25 nM) was activated with 10 μM iperoxo, and subsequently incubated for 60 min at room temperature with 1.5-fold molar access of bimane-labeled βarr1, 2-fold molar excess of Fab30 and SH3 (500 nM). Fluorescence emission spectra were collected in top-read mode, with excitation at 370 nm (16 nm bandpass) and emission scanning from 410 nm to 640 nm (10 nm bandpass) in 0.5 nm increments using a CLARIOstar microplate reader (BMG Labtech). Statistical comparisons were determined by comparing the area under the curves (*GraphPad Prism v9*) and peak fluorescence using one-way ANOVA with Dunnett’s post hoc test from three (M2V2R) or six (V2R membranes) independent experiments.

### FRET-based live-cell cAMP accumulation assay

cAMP production mediated by Gαs-coupled β2AR activation was measured using Epac2 (ICUE2) sensor containing a CFP and YFP FRET pair as described previously^31^. HEK293 cells stably expressing ICUE2 and β2AR were transfected with Src (2 ug) or the equivalent amount of control plasmids (pcDNA 3.1) with FuGENE®6 (Promega) transfection reagent according to the manufacturer’s instructions. 20 hours after transfection the cells were plated in poly-D-lysine-coated, black, clear-bottom 96-well plates (Corning) at a density of 50,000 cells per well. 24 hours after plating, cells were washed with PBS and incubated in imaging buffer (10 mM HEPES, 150 mM NaCl, 5 mM KCl, 1.5 mM MgCl2, 1.5 mM CaCl2, 10 mM glucose, 0.2% BSA, pH 7.4) for one hour at 37°C. The cells were stimulated with 10 μM isoproterenol (ISO), and FRET changes were measured in real-time using FlexStation 3 plate reader (Molecular Devices). The changes in the background-subtracted 480 nm/535 nm fluorescence emission ratio (CFP/YFP) are indicative of changes in cAMP levels. To quantify the overall cAMP response, the area under the curve (AUC) was calculated from the time-course data in *GraphPad Prism v9*.

### Data and statistical analysis

Data were analyzed using *GraphPad Prism v9*. Data represent the mean ±SEM of at least three independent experiments. Statistical significance for more than two groups were determined by one-way ANOVA with Dunnett’s post hoc test. Statistical significance for two groups was determined by Student’s t-test or Mann-Whitney test. No samples or data points were excluded from analysis.

## Data availability

The coordinates and the corresponding cryo-EM maps were deposited at the Protein Data Bank and the EMDB with the accession codes 9CX3 and EMD-45977 (SH3–βarr1-CC complex), 9CX9 and EMD-45982 (SH3–βarr1-N complex), 9BT8 and EMD-44881 (Src–βarr1-CC complex), respectively.

## Acknowledgements

We acknowledge the use of cryo-EM microscopes at the Shared Materials Instrumentation Facility (Duke University) and thank Nilakshee Bhattacharya for assistance with microscope operation. We are grateful to Liyin Huang for their helpful discussions throughout this work and to Yangyang Li for administrative assistance.

## Author contributions

N.P. and R.J.L. conceived the study. N.P., B.N.T., H.B., L.L., D.K.B., R.R.A., J.K., A.W.K., B.P., K.X., S.L., R.O., S.A., and X.Z. performed the experiments. N.P., B.N.T., H.B., D.K.B., R.R.A., J.K., A.W.K., B.P., and K.X. analyzed the data. N.P. wrote the paper with input from all authors. A.G. and R.J.L. supervised the work.

## Correspondence and requests for materials

The materials presented are available upon request from Robert J. Lefkowitz.

## Supplementary Information

**Supplementary Fig. 1.**
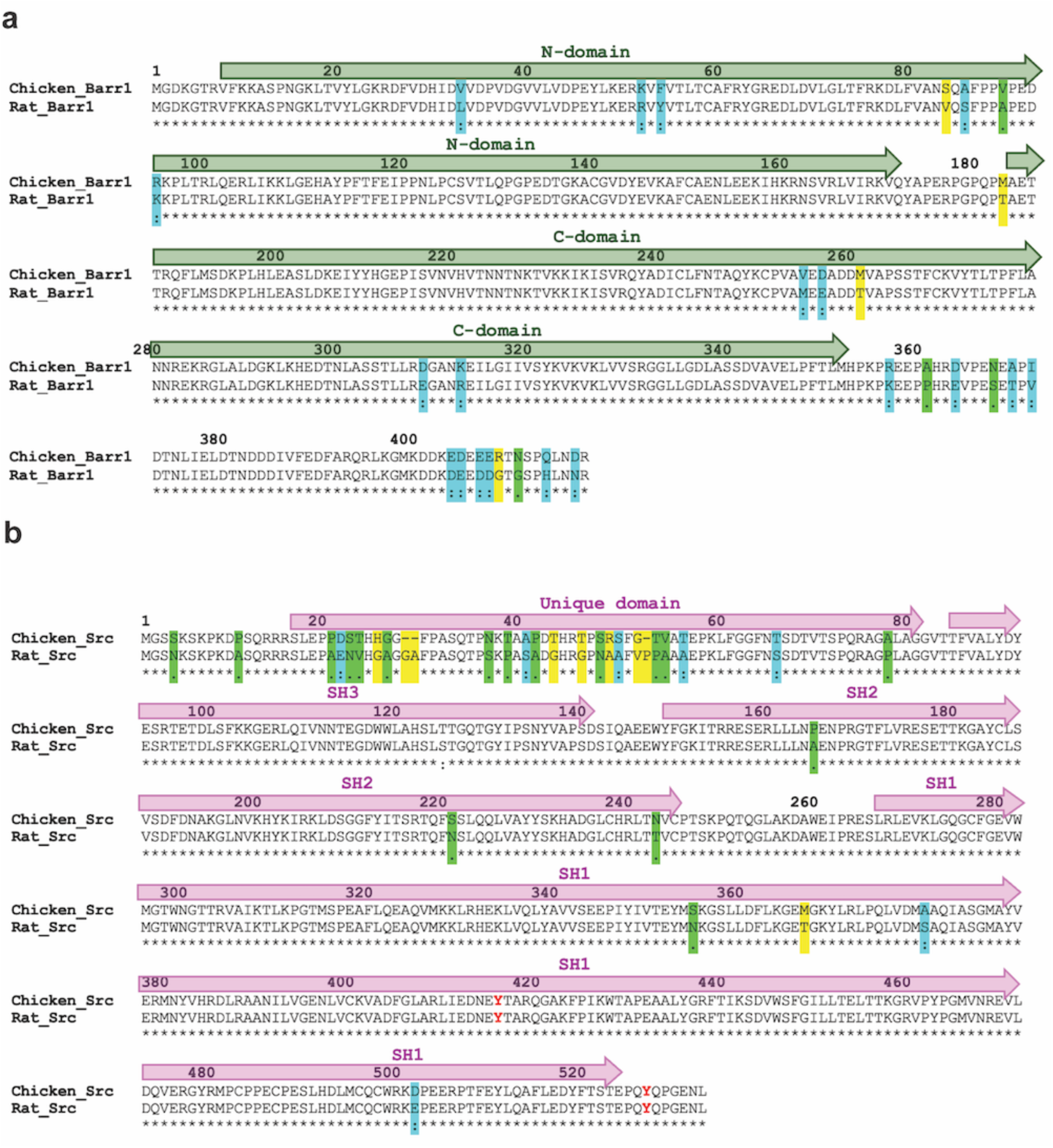
Alignment of rat (*Rattus norvegicus*) and chicken (*Gallus gallus*) sequences of βarr1 (a) and Src (b). Fully conserved residues are marked with asterisk (*); strongly conserved residues are marked with colon (:) and shaded in cyan; weakly conserved residues are marked with period (.) and shaded in green; non-conserved residues are shaded in yellow. Src active loop tyrosine (Y416) and C-tail tyrosine (Y527) are colored in red. Domains are indicated with arrows and labeled. Sequence alignment was performed in Clustal Omega (https://www.ebi.ac.uk/jdispatcher/msa/clustalo).

**Supplementary Fig. 2.**
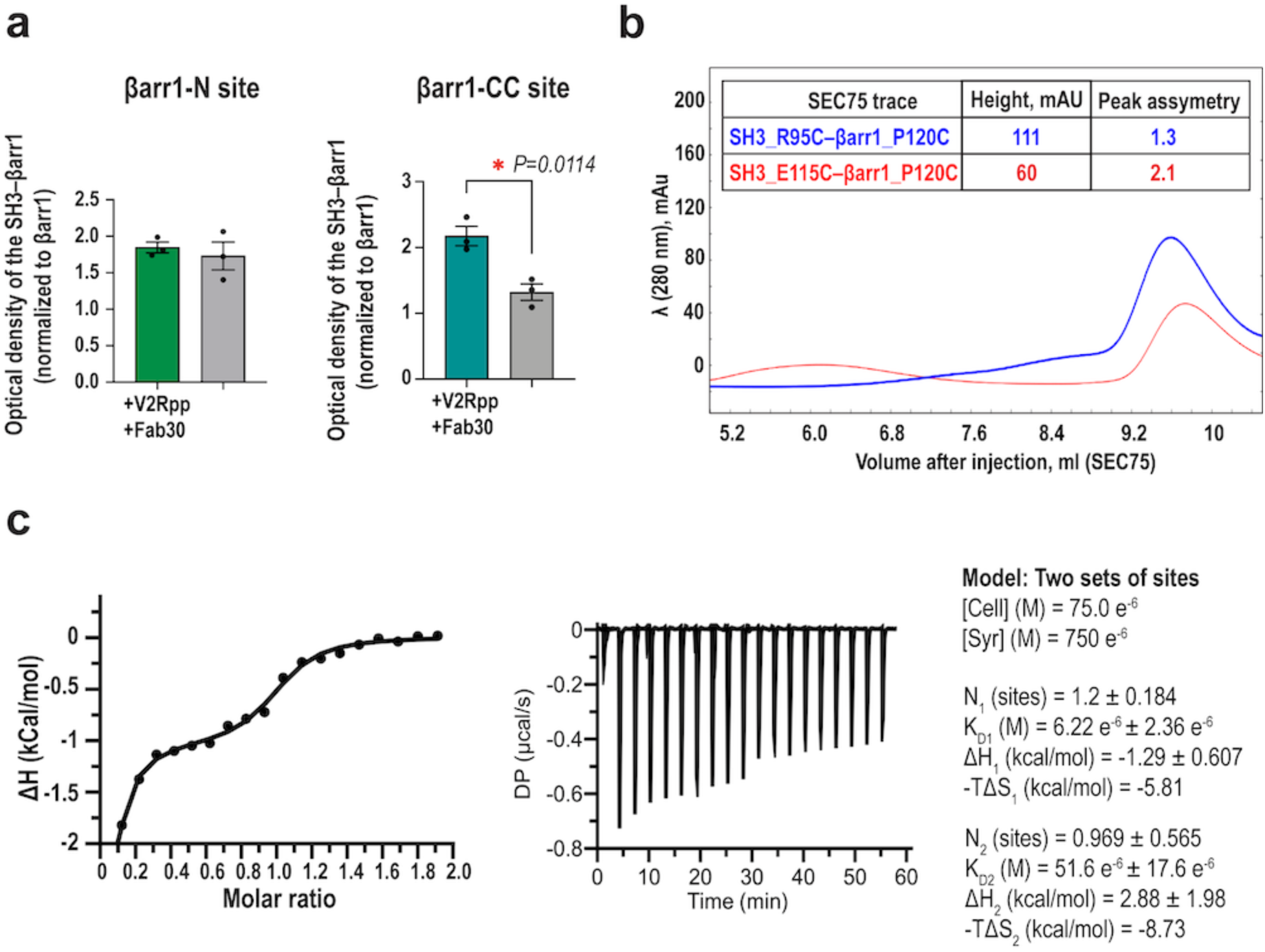
βarr1 uses two distinct sites to bind SH3. **a,** Densitometry analysis of SH3–βarr1 disulfide trapping with and without V2Rpp (synthetic phosphopeptide mimicking the C-tail of vasopressin 2 receptor) and stabilizing antibody Fab30 (mean ± standard error of mean (SEM), n=3). Statistical analysis was performed using Student’s t-test. **b,** Analytical size-exclusion chromatography profile of SH3_R95C–βarr1_P120C and SH3_E115C–βarr1_P120C disulfide trapped complexes. **c,** Isothermal titration calorimetry of SH3 binding to βarr1–V2Rpp. Top panel: integrated heat (after deducting heat of dilution) per injection of SH3 into βarr1–V2Rpp solution in the cell based on the molar ratio of each injection. Bottom panel: raw titration data showing the heat rate associated with each dilution per injection versus time into βarr1–V2Rpp solution in the cell. Right panel: thermodynamic parameters of SH3–βarr1–V2Rpp. The data are representative of three independent experiments.

**Supplementary Fig. 3.**
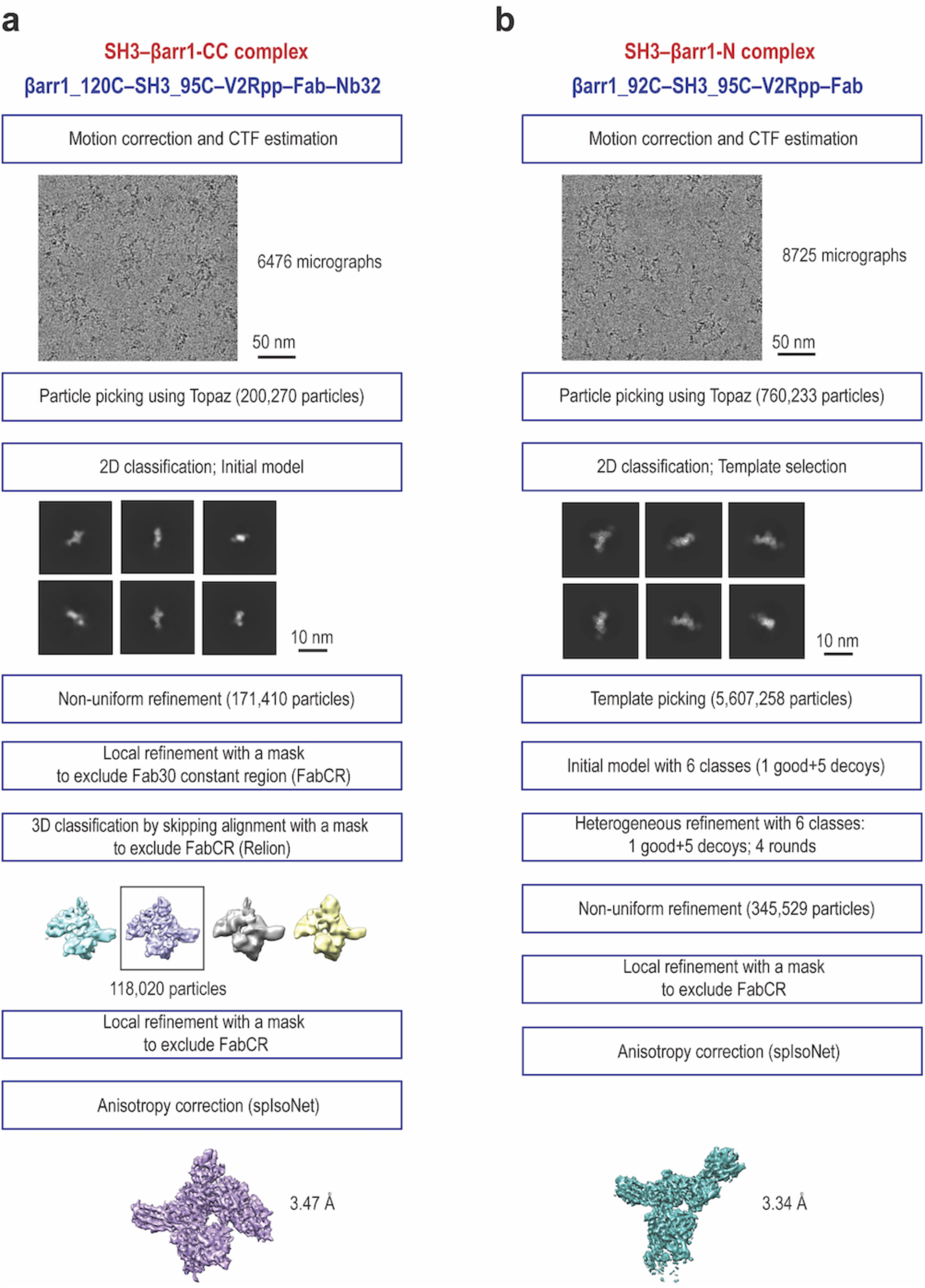
Flow chart of cryo-EM data processing of SH3–βarr1-CC (a) and SH3–βarr1-N (b) complexes.

**Supplementary Fig. 4.**
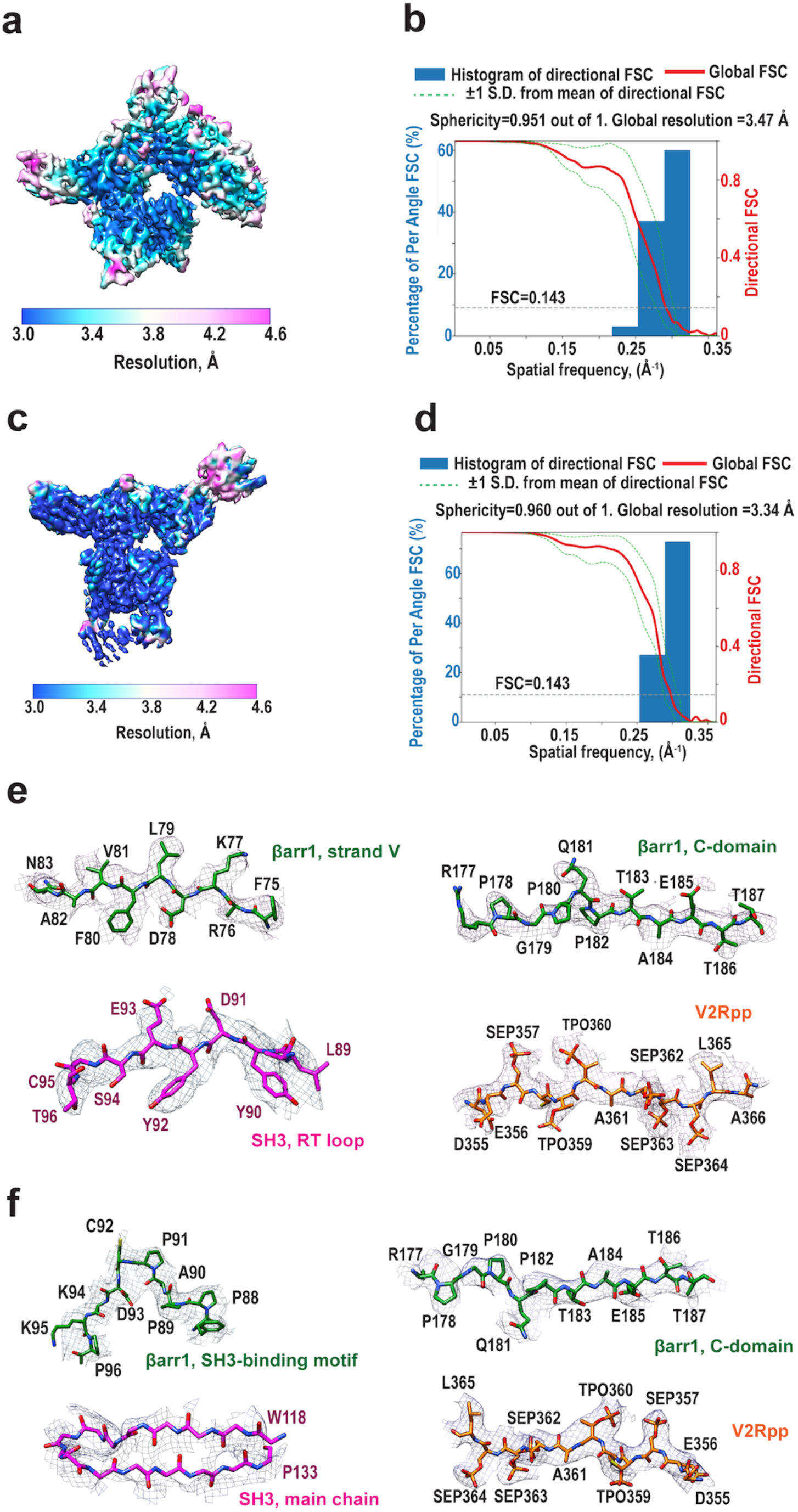
Cryo-electron microscopy of SH3–βarr1 complexes. **a-c,** Cryo-EM maps of SH3–βarr1-CC (**a**) and SH3–βarr1-N (**c**) coloured by local resolution; resolution is reported at the Fourier Shell Correlation (FSC) threshold of 0.143. Map contour level is 0.83 for SH3–βarr1-CC (**a**) and 0.50 for SH3–βarr1-N (**c**). **b-d,** Histogram and directional FSC plot measuring directional resolution anisotropy for SH3–βarr1-CC (**b**) and 0.50 for SH3–βarr1-N (**d**). Sphericity values were determined at an FSC threshold of 0.5 with the 3DFSC software.^73^ **e-f,** Cryo-EM density at different parts of the SH3–βarr1-CC (**e**) and SH3–βarr1-N (**f**) complexes. The upsample maps are used (0.69 Å/pix); map contour level is 0.3 (SH3–βarr1-CC) and 0.2 (SH3–βarr1-N).

**Supplementary Fig. 5.**
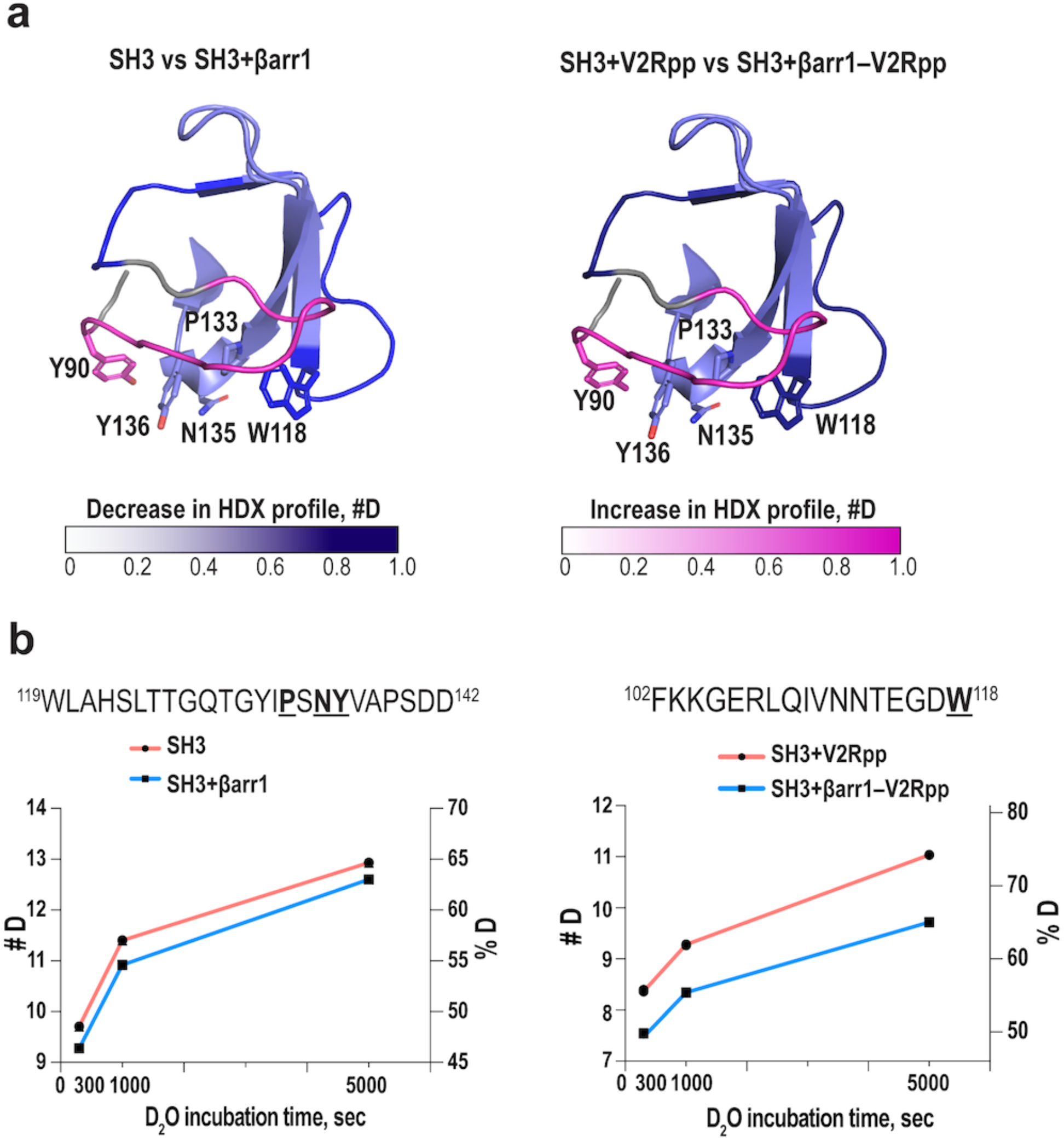
HDX profile changes in SH3 upon co-incubation with βarr1. **a,** Structure of SH3 (PDB: 2PTK) upon co-incubation with free βarr1 (left panel) or V2Rpp-activated βarr1 (right panel). Regions with decreased and increased deuterium uptake are shaded in dark-blue and magenta, respectively. Only regions that showed differences between the states in deuterium uptake (>0.2 #D) in at least two overlapping peptides are indicated. **b,** The HDX profile of representative peptides in SH3. The residues interacting with βarr1 in the structures are shown in bold and underlined.

**Supplementary Fig. 6.**
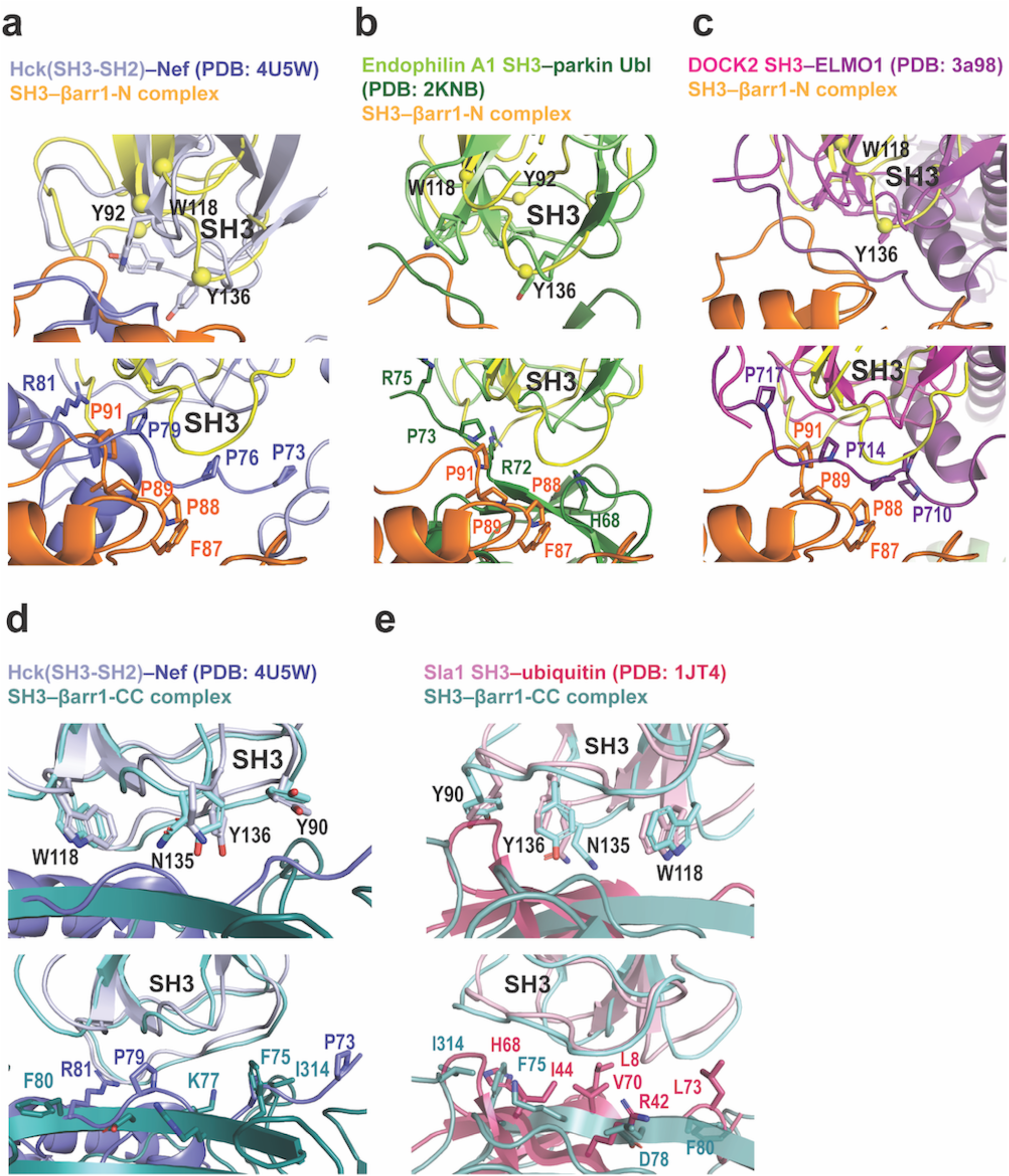
Structural superpositions of SH3 domains in SH3–βarr1-N (yellow), SH3–βarr1-CC (cyan) and other SH3-binding complexes. **a,** SH3–βarr1-N (yellow) and Hck(SH3-SH2)-Nef (slate). **b,** SH3–βarr1-N (yellow) and endophilin A1 SH3–parkin Ubl (green). **c,** SH3–βarr1-N (yellow) and DOCK2 SH3–ELMO 1 (magenta). **d,** SH3–βarr1-CC (cyan) and Hck(SH3-SH2)-Nef (slate). **e,** SH3–βarr1-CC (cyan) and Sla1 SH3-ubiquitin (pink). The top panel shows the interacting residues in SH3 (Src SH3 numbering is used), the bottom panel shows interacting residues in βarr1 and the SH3-binding protein.

**Supplementary Fig. 7.**
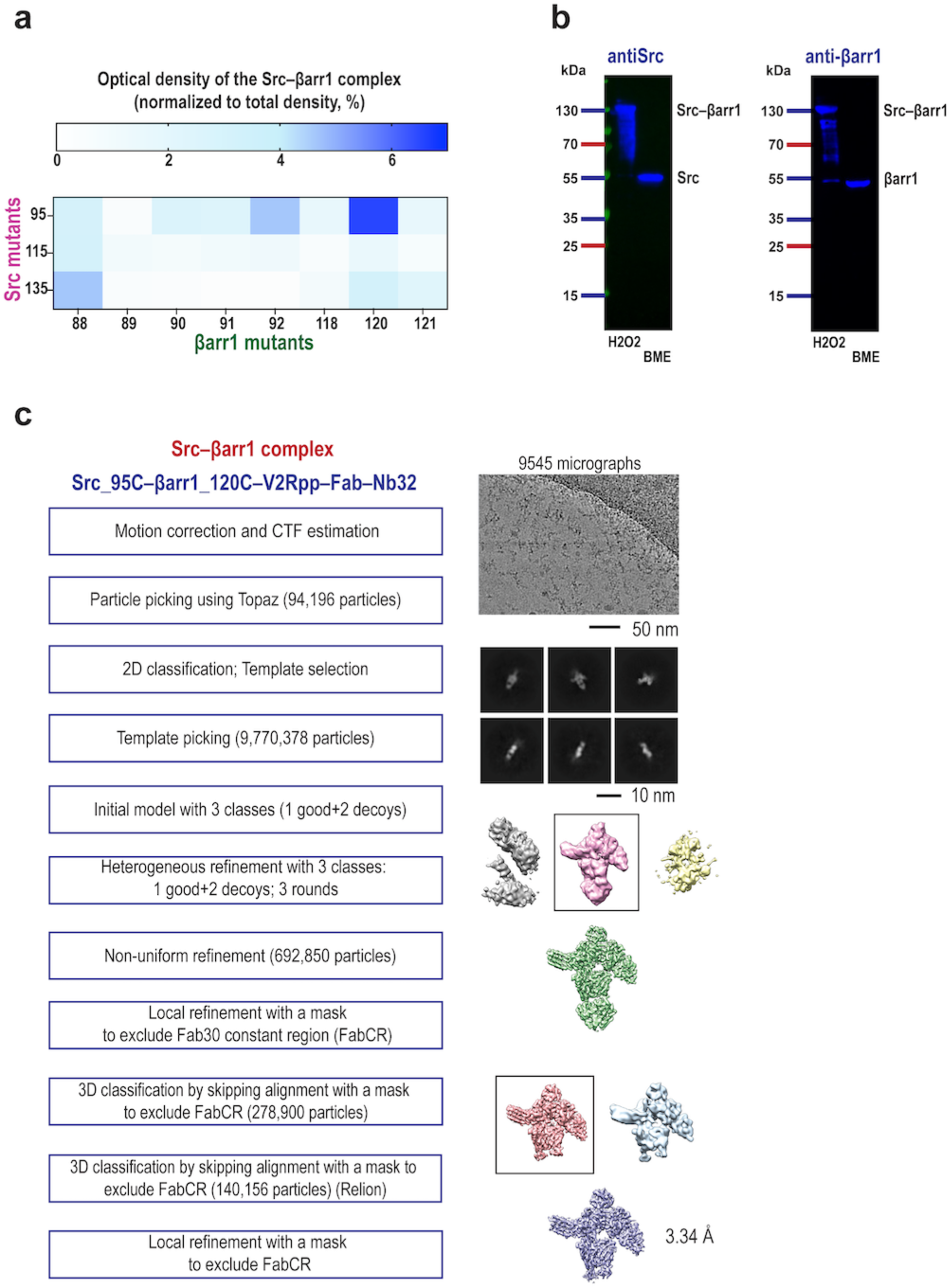
Complex formation and cryo-EM data processing of Src–βarr1-CC. **a,** Complex formation between Src and βarr1 mutants revealed by disulfide trapping; densitometry analysis of Coomassie blue gels. The Src–βarr1-CC complex band was normalized to the total density of all bands in each sample (mean, n=3-9). Prior to disulfide trapping reactions, βarr1 was activated by V2Rpp. **b,** Western blot of disulfide trapping reaction with βarr1_120C and Src_R95C. 130-kDa band is detected by both anti-βarr1 (A1CT) and anti-Src antibodies, confirming that the band is the covalent Src–βarr1-CC complex. **c,** Flow chart of cryo-EM data processing of the Src–βarr1-CC complex.

**Supplementary Fig. 8.**
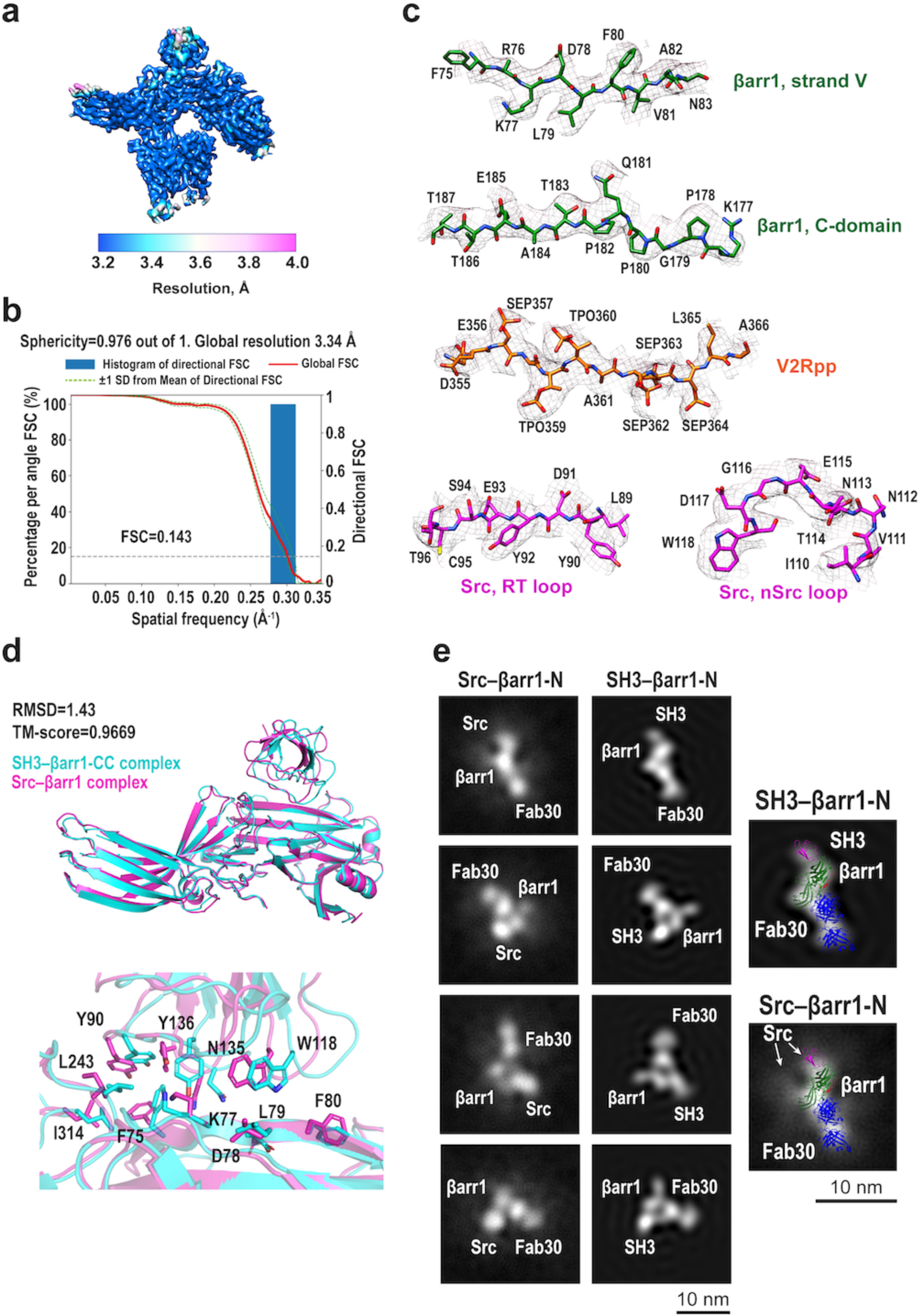
Cryo-electron microscopy of Src–βarr1 complexes. **a,** Cryo-EM map of the Src–βarr1-CC complex coloured by local resolution; resolution is reported at the FSC threshold of 0.143. Map contour level is 0.55. **b,** Histogram and directional FSC plot. A sphericity of 0.976 determined at an FSC threshold of 0.5 indicates isotropic angular distribution. Directional FSC determination was performed with the 3DFSC software.^73^ **c,** Cryo-EM density at different parts of Src–βarr1-CC map. The upsample map is used (0.72 Å/pix); map contour level is 0.1. **d,** The structural superposition of the Src–βarr1-CC and SH3–βarr1-CC complexes (Fab30, V2Rpp and Nb32 are not shown). RMSD and the TM-score of the structures was calculated using the TM-score function^74^. **e,** Matching of the 2D classes of the Src–βarr1-N complex with the projections of the SH3–βarr1-N map low-pass filtered to 25 Å using cluster selection mode in Reference Based Auto Select 2D module in *CryoSPARC* (left panel) and representative 2D classes of the complexes with the fitted model of SH3–βarr1-N (right panel).

**Supplementary Fig. 9.**
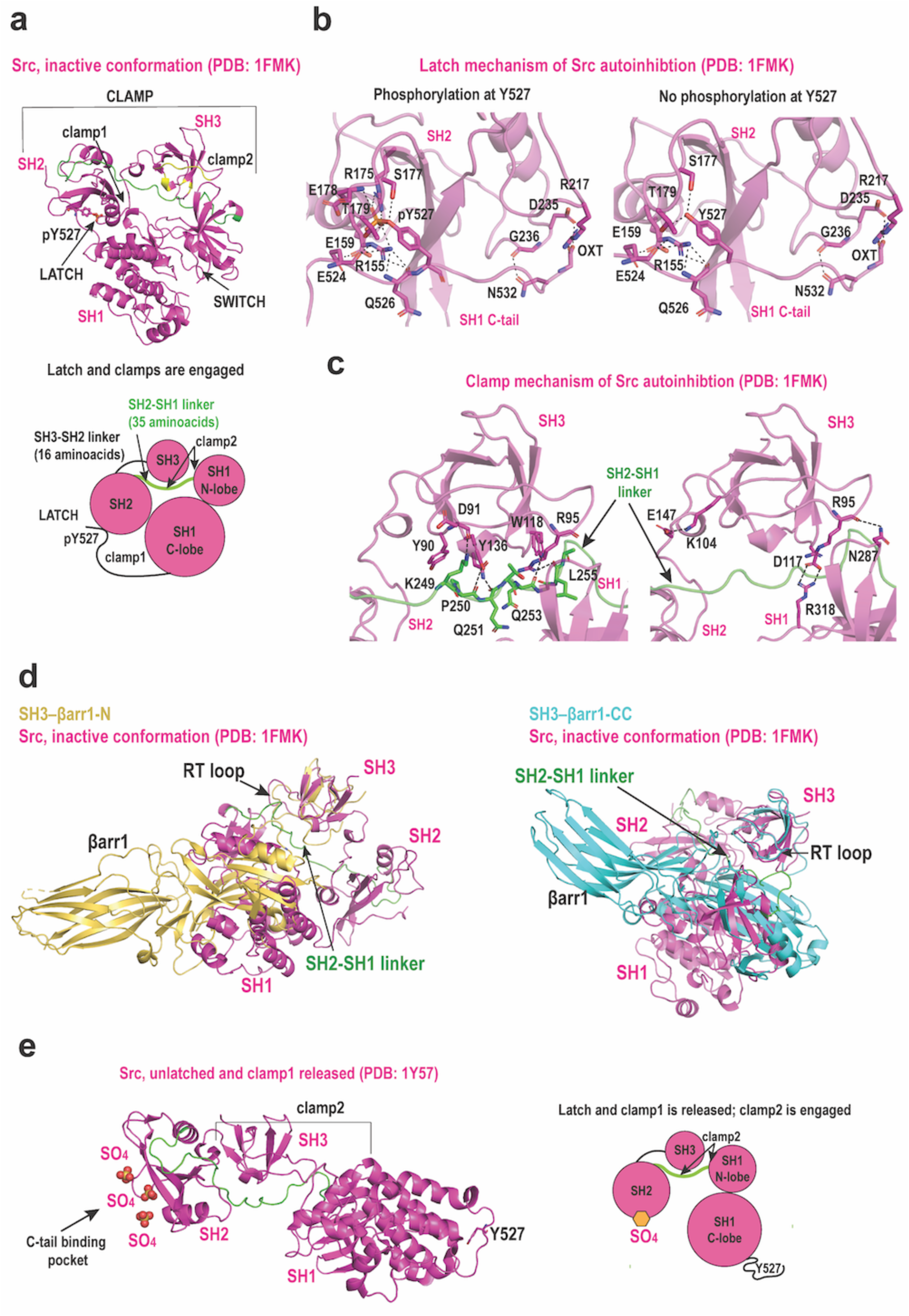
Autoinhibition of Src. **a,** Structure of autoinhibited Src (PDB: 1FMK), magenta. Parts of SH3 interacting with βarr1 are colored in yellow. SH2-SH1 linker is colored in green; phosphorylated Y527 is labeled. **b-c,** Latch (**b**) and clamp (**c**) mechanisms of Src autoinhibition (PDB: 1FMK), cartoon representation (magenta). The hydrogen bonds are depicted as dashed lines; interacting residues are labeled. **d,** Structural superpositions of SH3–βarr1-N (yellow) and SH3–βarr1-CC (cyan) with inactive Src (PDB: 1FMK, magenta). **e,** Structure of unphosphorylated Src (PDB: 1Y57), magenta. SH2-SH1 linker is colored in green; phosphorylated Y527 is labeled. Note that the C-tail binding pocket on SH2 is occupied by sulfate ions originating from crystallization solution.

**Supplementary Table 1.** Cryo-EM data collection, refinement and validation statistics.

|  | **SH3_95C-βarr1_120C-**<br>**V2Rpp-Fab30-Nb32**<br>**(SH3-βarr1-CC)**<br>**(EMD-45977)**<br>**(PDB 9CX3)** | **SH3_95C-βarr1_92C-**<br>**V2Rpp-Fab30**<br>**(SH3-βarr1-N)**<br>**(EMD-45982)**<br>**(PDB 9CX9)** | **Src_95C-βarr1_120C-**<br>**V2Rpp-Fab30-Nb32**<br>**(Src-βarr1-CC)**<br>**(EMD-44881)**<br>**(PDB 9BT8)** |
| --- | --- | --- | --- |
| **Data collection and processing** |  |  |  |
| Magnification | 81,000 | 81,000 | 81,000 |
| Voltage (kV) | 300 | 300 | 300 |
| Electron exposure (e-/$\text{\AA}^2$) | 58.5 | 53.8 | 54.6 |
| Defocus range ($\mu\text{m}$) | -0.8 to -2.5 | -0.8 to -2.5 | -0.8 to -2.5 |
| Pixel size ($\text{\AA}$) | 1.08 (collection)<br>1.3824 (final) | 1.08 (collection)<br>1.3824 (final) | 1.08 (collection)<br>1.44 (final) |
| Symmetry imposed | C1 | C1 | C1 |
| Initial particle projections (no.) | 200,270 | 5,607,258 | 9,770,378 |
| Final particle projections (no.) | 118,020 | 345,529 | 140,156 |
| Map resolution ($\text{\AA}$) | 3.47 | 3.34 | 3.32 |
| FSC threshold | 0.143 | 0.143 | 0.143 |
| Map resolution range ($\text{\AA}$) | 3.10-6.97 | 3.05-6.90 | 3.23-11.33 |
| **Refinement** |  |  |  |
| Initial model used (PDB code) | 6NI2 | 4JQI | 8U7A |
| Model resolution ($\text{\AA}$) | 3.47 | 3.3 | 3.2 |
| FSC threshold | 0.143 | 0.143 | 0.143 |
| Map sharpening $B$ factor ($\text{\AA}^2$) | -80.3 | -83.2 | -77.5 |
| **Model composition** |  |  |  |
| Non-hydrogen atoms | 5627 | 4580 | 5713 |
| Protein residues | 720 | 609 | 728 |
| Ligands | 0 | 0 | 0 |
| **$B$ factors ($\text{\AA}^2$)** |  |  |  |
| Protein (min/max/mean) | 20.22/103.43/47.54 | 8.88/113.35/37.29 | 10.51/115.11/53.64 |
| Ligand | --- | --- |  |
| **R.m.s. deviations** |  |  |  |
| Bond lengths ($\text{\AA}$) | 0.005 | 0.004 | 0.004 |
| Bond angles ($^\circ$) | 0.709 | 0.988 | 0.572 |
| **Validation** |  |  |  |
| MolProbity score | 1.93 | 1.56 | 1.55 |
| Clashscore | 11.22 | 7.59 | 6.88 |
| Poor rotamers (%) | 0.16 | 0.21 | 0.32 |
| **Ramachandran plot** |  |  |  |
| Favored (%) | 94.78 | 97.25 | 97.00 |
| Allowed (%) | 5.22 | 2.75 | 3.00 |
| Disallowed (%) | 0.00 | 0.00 | 0.00 |

**Supplementary Video 1.** SH3–βarr1-CC complex, cryo-EM map and model colored by subunit (green, βarr1; magenta, SH3; red, V2Rpp; blue, Fab30; yellow, Nb32). Interacting residues are shown as sticks, hydrogen bonds are indicated as dashed lines. Map contour level is 0.75.

**Supplementary Video 2.** SH3–βarr1-N complex, cryo-EM map and model colored by subunit (green, βarr1; magenta, SH3; red, V2Rpp; blue, Fab30). Map contour level is 0.52.

**Supplementary Video 3.** βarr1 activates Src by SH3 domain displacement. Inactive Src is constrained by the intramolecular interactions between SH3 (magenta), SH2 (cyan) and SH1 (blue). βarr1 (green, the βarr1-CC site is shown) binds the aromatic surface of SH3 and displaces the SH2-SH1 linker (yellow) releasing the catalytic (SH1) domain of Src. Transparent coloring indicates flexible regions, including the inducible snap lock, C-terminal tail, and liberated SH1 domain. The displaced SH2-SH1 linker is shown as dashed line.

## REFERENCES

1. Lefkowitz, R.J. Arrestins come of age: a personal historical perspective. Prog Mol Biol Transl Sci 118, 3–18 (2013).

2. Peterson, Y.K. & Luttrell, L.M. The Diverse Roles of Arrestin Scaffolds in G Protein-Coupled Receptor Signaling. Pharmacol Rev 69, 256–297 (2017).

3. Yang, F. et al. Allosteric mechanisms underlie GPCR signaling to SH3-domain proteins through arrestin. Nature Chemical Biology 14, 876-+ (2018).

4. Pakharukova, N., Masoudi, A., Pani, B., Staus, D.P. & Lefkowitz, R.J. Allosteric activation of proto-oncogene kinase Src by GPCR-beta-arrestin complexes. J Biol Chem 295, 16773–16784 (2020).

5. Zang, Y., Kahsai, A.W., Pakharukova, N., Huang, L.Y. & Lefkowitz, R.J. The GPCR-beta-arrestin complex allosterically activates C-Raf by binding its amino terminus. J Biol Chem 297, 101369 (2021).

6. Kahsai, A.W. et al. Signal transduction at GPCRs: Allosteric activation of the ERK MAPK by β-arrestin. Proceedings of the National Academy of Sciences of the United States of America 120(2023).

7. Karkkainen, S. et al. Identification of preferred protein interactions by phage-display of the human Src homology-3 proteome. EMBO Rep 7, 186–91 (2006).

8. Li, S.S. Specificity and versatility of SH3 and other proline-recognition domains: structural basis and implications for cellular signal transduction. Biochem J 390, 641–53 (2005).

9. Luttrell, L.M. et al. Beta-arrestin-dependent formation of beta2 adrenergic receptor-Src protein kinase complexes. Science 283, 655–61 (1999).

10. Xiao, K. et al. Revealing the architecture of protein complexes by an orthogonal approach combining HDXMS, CXMS, and disulfide trapping. Nat Protoc 13, 1403–1428 (2018).

11. Cahill, T.J., 3rd et al. Distinct conformations of GPCR-beta-arrestin complexes mediate desensitization, signaling, and endocytosis. Proc Natl Acad Sci U S A 114, 2562–2567 (2017).

12. Shukla, A.K. et al. Structure of active beta-arrestin-1 bound to a G-protein-coupled receptor phosphopeptide. Nature 497, 137–41 (2013).

13. Lee, C.H., Saksela, K., Mirza, U.A., Chait, B.T. & Kuriyan, J. Crystal structure of the conserved core of HIV-1 Nef complexed with a Src family SH3 domain. Cell 85, 931–42 (1996).

14. Alvarado, J.J., Tarafdar, S., Yeh, J.I. & Smithgall, T.E. Interaction with the Src homology (SH3-SH2) region of the Src-family kinase Hck structures the HIV-1 Nef dimer for kinase activation and effector recruitment. J Biol Chem 289, 28539–53 (2014).

15. He, Y., Hicke, L. & Radhakrishnan, I. Structural basis for ubiquitin recognition by SH3 domains. J Mol Biol 373, 190–6 (2007).

16. Trempe, J.F. et al. SH3 domains from a subset of BAR proteins define a Ubl-binding domain and implicate parkin in synaptic ubiquitination. Mol Cell 36, 1034–47 (2009).

17. Hanawa-Suetsugu, K. et al. Structural basis for mutual relief of the Rac guanine nucleotide exchange factor DOCK2 and its partner ELMO1 from their autoinhibited forms. Proceedings of the National Academy of Sciences of the United States of America 109, 3305–3310 (2012).

18. Teyra, J. et al. Comprehensive Analysis of the Human SH3 Domain Family Reveals a Wide Variety of Non-canonical Specificities. Structure 25, 1598–1610 e3 (2017).

19. Saksela, K. & Permi, P. SH3 domain ligand binding: What’s the consensus and where’s the specificity? FEBS Lett 586, 2609–14 (2012).

20. Harrison, S.C. Variation on an Src-like theme. Cell 112, 737–40 (2003).

21. Xu, W., Harrison, S.C. & Eck, M.J. Three-dimensional structure of the tyrosine kinase c-Src. Nature 385, 595–602 (1997).

22. Porter, M., Schindler, T., Kuriyan, J. & Miller, W.T. Reciprocal regulation of Hck activity by phosphorylation of Tyr(527) and Tyr(416). Effect of introducing a high affinity intramolecular SH2 ligand. J Biol Chem 275, 2721–6 (2000).

23. Young, M.A., Gonfloni, S., Superti-Furga, G., Roux, B. & Kuriyan, J. Dynamic coupling between the SH2 and SH3 domains of c-Src and hck underlies their inactivation by C-terminal tyrosine phosphorylation. Cell 105, 115–126 (2001).

24. Bous, J. et al. Structure of the vasopressin hormone-V2 receptor-beta-arrestin1 ternary complex. Sci Adv 8, eabo7761 (2022).

25. Staus, D.P. et al. Structure of the M2 muscarinic receptor-beta-arrestin complex in a lipid nanodisc. Nature 579, 297–302 (2020).

26. Huang, W. et al. Structure of the neurotensin receptor 1 in complex with beta-arrestin 1. Nature 579, 303–308 (2020).

27. Lee, Y. et al. Molecular basis of beta-arrestin coupling to formoterol-bound beta(1)- adrenoceptor. Nature 583, 862-+ (2020).

28. O’Hayre, M. et al. Genetic evidence that β-arrestins are dispensable for the initiation of β-adrenergic receptor signaling to ERK. Science Signaling 10 (2017).

29. Stramiello, M. & Wagner, J.J. D1/5 receptor-mediated enhancement of LTP requires PKA, Src family kinases, and NR2B-containing NMDARs. Neuropharmacology 55, 871–877 (2008).

30. Kaya, A.I., Perry, N.A., Gurevich, V.V. & Iverson, T.M. Phosphorylation barcode-dependent signal bias of the dopamine D1 receptor. Proc Natl Acad Sci U S A 117, 14139–14149 (2020).

31. Violin, J.D. et al. β-adrenergic receptor signaling and desensitization elucidated by quantitative modeling of real time cAMP dynamics. Journal of Biological Chemistry 283, 2949–2961 (2008).

32. Qin, L., et al. Structural biology. Crystal structure of the chemokine receptor CXCR4 in complex with a viral chemokine. Science 347, 1117–22 (2015).

33. Rosenbaum, D.M. et al. Structure and function of an irreversible agonist-beta(2) adrenoceptor complex. Nature 469, 236–40 (2011).

34. Chen, Q.Y. et al. Structures of rhodopsin in complex with G-protein-coupled receptor kinase 1. Nature 595, 600-+ (2021).

35. Scott, J.D. & Pawson, T. Cell Signaling in Space and Time: Where Proteins Come Together and When They’re Apart. Science 326, 1220–1224 (2009).

36. Moarefi, I. et al. Activation of the Src-family tyrosine kinase Hck by SH3 domain displacement. Nature 385, 650–3 (1997).

37. Alexandropoulos, K. & Baltimore, D. Coordinate activation of c-Src by SH3-and SH2-binding sites on a novel, p130(Cas)-related protein, Sin. Genes & Development 10, 1341–1355 (1996).

38. Kypta, R.M., Goldberg, Y., Ulug, E.T. & Courtneidge, S.A. Association between the Pdgf Receptor and Members of the Src Family of Tyrosine Kinases. Cell 62, 481–492 (1990).

39. Cobb, B.S., Schaller, M.D., Leu, T.H. & Parsons, J.T. Stable Association of Pp60(Src) and Pp59(Fyn) with the Focal Adhesion-Associated Protein-Tyrosine Kinase, Pp125(Fak). Molecular and Cellular Biology 14, 147–155 (1994).

40. Burnham, M.R. et al. Regulation of c-SRC activity and function by the adapter protein CAS. Molecular and Cellular Biology 20, 5865–5878 (2000).

41. Bjorge, J.D., Jakymiw, A. & Fujita, D.J. Selected glimpses into the activation and function of Src kinase. Oncogene 19, 5620–5635 (2000).

42. Roskoski, R. Src protein-tyrosine kinase structure, mechanism, and small molecule inhibitors. Pharmacological Research 94, 9–25 (2015).

43. Mitra, S.K. & Schlaepfer, D.D. Integrin-regulated FAK-Src signaling in normal and cancer cells. Curr Opin Cell Biol 18, 516–23 (2006).

44. Thomsen, A.R.B. et al. GPCR-G Protein-beta-Arrestin Super-Complex Mediates Sustained G Protein Signaling. Cell 166, 907–919 (2016).

45. Nguyen, A.H. et al. Structure of an endosomal signaling GPCR-G protein-β-arrestin megacomplex. Nature Structural & Molecular Biology 26, 1123-+ (2019).

46. Frame, M.C. Src in cancer: deregulation and consequences for cell behaviour. Biochim Biophys Acta 1602, 114–30 (2002).

47. Benovic, J.L. et al. Functional desensitization of the isolated beta-adrenergic receptor by the beta-adrenergic receptor kinase: potential role of an analog of the retinal protein arrestin (48-kDa protein). Proc Natl Acad Sci U S A 84, 8879–82 (1987).

48. Lohse, M.J., Benovic, J.L., Codina, J., Caron, M.G. & Lefkowitz, R.J. Beta-Arrestin - a Protein That Regulates Beta-Adrenergic-Receptor Function. Science 248, 1547–1550 (1990).

49. Gutkind, J.S. & Kostenis, E. Arrestins as rheostats of GPCR signalling. Nature Reviews Molecular Cell Biology 19, 615–616 (2018).

50. Staus, D.P. et al. Sortase ligation enables homogeneous GPCR phosphorylation to reveal diversity in beta-arrestin coupling. Proc Natl Acad Sci U S A 115, 3834–3839 (2018).

51. Paduch, M. et al. Generating conformation-specific synthetic antibodies to trap proteins in selected functional states. Methods 60, 3–14 (2013).

52. Shukla, A.K. et al. Visualization of arrestin recruitment by a G-protein-coupled receptor. Nature 512, 218–222 (2014).

53. Seeliger, M.A. et al. High yield bacterial expression of active c-Abl and c-Src tyrosine kinases. Protein Sci 14, 3135–9 (2005).

54. Nobles, K.N., Guan, Z., Xiao, K., Oas, T.G. & Lefkowitz, R.J. The active conformation of beta-arrestin1: direct evidence for the phosphate sensor in the N-domain and conformational differences in the active states of beta-arrestins1 and -2. J Biol Chem 282, 21370–81 (2007).

55. Rizk, S.S. et al. Allosteric control of ligand-binding affinity using engineered conformation-specific effector proteins. Nat Struct Mol Biol 18, 437–42 (2011).

56. Attramadal, H. et al. Beta-Arrestin2, a Novel Member of the Arrestin Beta-Arrestin Gene Family. Journal of Biological Chemistry 267, 17882–17890 (1992).

57. Chen, S., Li, J., Vinothkumar, K.R. & Henderson, R. Interaction of human erythrocyte catalase with air-water interface in cryoEM. Microscopy (Oxf) 71, i51–i59 (2022).

58. Punjani, A., Rubinstein, J.L., Fleet, D.J. & Brubaker, M.A. cryoSPARC: algorithms for rapid unsupervised cryo-EM structure determination. Nat Methods 14, 290–296 (2017).

59. Bepler, T. et al. Positive-unlabeled convolutional neural networks for particle picking in cryo-electron micrographs. Nature Methods 16, 1153-+ (2019).

60. Scheres, S.H. RELION: implementation of a Bayesian approach to cryo-EM structure determination. J Struct Biol 180, 519–30 (2012).

61. Liu, Y.T., Fan, H., Hu, J.J. & Zhou, Z.H. Overcoming the preferred-orientation problem in cryo-EM with self-supervised deep learning. Nat Methods 22, 113–123 (2025).

62. Sanchez-Garcia, R. et al. DeepEMhancer: a deep learning solution for cryo-EM volume post-processing. Commun Biol 4, 874 (2021).

63. Pettersen, E.F. et al. UCSF Chimera--a visualization system for exploratory research and analysis. J Comput Chem 25, 1605–12 (2004).

64. Emsley, P., Lohkamp, B., Scott, W.G. & Cowtan, K. Features and development of Coot. Acta Crystallogr D Biol Crystallogr 66, 486–501 (2010).

65. Croll, T.I. ISOLDE: a physically realistic environment for model building into low-resolution electron-density maps. Acta Crystallographica Section D-Structural Biology 74, 519–530 (2018).

66. Pettersen, E.F. et al. UCSF ChimeraX: Structure visualization for researchers, educators, and developers. Protein Sci 30, 70–82 (2021).

67. Adams, P.D. et al. PHENIX: building new software for automated crystallographic structure determination. Acta Crystallogr D Biol Crystallogr 58, 1948–54 (2002).

68. Kim, D.N. et al. Cryo_fit: Democratization of flexible fitting for cryo-EM. J Struct Biol 208, 1–6 (2019).

69. Kidmose, R.T. et al. Namdinator - automatic molecular dynamics flexible fitting of structural models into cryo-EM and crystallography experimental maps. Iucrj 6, 526–531 (2019).

70. Chen, V.B. et al. MolProbity: all-atom structure validation for macromolecular crystallography. Acta Crystallogr D Biol Crystallogr 66, 12–21 (2010).

71. Barker, S.C. et al. Characterization of pp60c-src tyrosine kinase activities using a continuous assay: autoactivation of the enzyme is an intermolecular autophosphorylation process. Biochemistry 34, 14843–51 (1995).

72. Strachan, R.T. et al. Divergent Transducer-specific Molecular Efficacies Generate Biased Agonism at a G Protein-coupled Receptor (GPCR). Journal of Biological Chemistry 289, 14211–14224 (2014).

73. Tan, Y.Z. et al. Addressing preferred specimen orientation in single-particle cryo-EM through tilting. Nature Methods 14, 793-+ (2017).

74. Zhang, Y. & Skolnick, J. Scoring function for automated assessment of protein structure template quality. Proteins 57, 702–10 (2004).

